# Corrections in single cell migration path in vivo are controlled by pulses in polar Rac1 activation

**DOI:** 10.1101/2025.02.24.639819

**Authors:** Dennis Hoffmann, Tal Agranov, Lucas Kühl, Benjamin D. Simons, Nir Gov, Erez Raz

**Author notes:** Correspondence (T.A.), (E.R.).

## Abstract

Directed migration of single cells is central to a large number of processes in development and adult life. Corrections to the migration path of cells are often characterized by periodic loss of polarity that is followed by the generation of a new leading edge in response to guidance cues, a behavior termed ‘run and tumble’. While this phenomenon is essential for accurate arrival at migration targets, the precise molecular mechanisms responsible for the periodic changes in cell polarity are unknown. To investigate this issue, we employ germ cells in live zebrafish embryos as an *in vivo* model and show that a tunable molecular network controls periodic pulsations of Rac1 activity and actin polymerization. This process, which we term ‘polar pulsations’, is responsible for the transitions between the run and tumble phases. In addition, we provide evidence for the role of apolar blebbing activity during tumble phases in erasing the memory of the previous front-back polarity of the migrating cell. To understand how the molecular components give rise to this distinct behavior, we develop a minimal mathematical model of the biochemical network that accounts for the observed cell behavior. Together, our *in vivo* findings and the mathematical model suggest that a pulsatory signaling network regulates the accuracy of individual cell migration.

## Introduction

Migration of cells is a key process in numerous developmental and physiological processes.^1–4^ Consequently, abnormal migration of cells is associated with defects in embryonic development,^5^ organogenesis,^6,7^ wound healing,^8^ and immune responses,^9^ and also leads to cancer metastasis.^10^ Motile cells are guided toward their target by extracellular signals that act as chemotactic attractive or repulsive cues.^11^ Interestingly, changes in the direction of migration often involve a phase in which the cells stop migrating, after which they re-establish motility. This “run and tumble” behavior was first described in bacteria,^12^ but similar migration patterns are found in other cell types, such as neutrophils,^13^ T cells,^14^ microglia,^15^ axon growth cones,^16^ and dendritic cells,^17^ underscoring the general importance of the behavior. Zebrafish primordial germ cells (PGCs) also display run behavior, during which they form protrusions in the direction of movement, and tumble phases when the cells lose morphological polarity, stop migrating, and extend protrusions in all directions.^18^ Importantly, the periodic acquisition and loss of morphological polarity, which is important for the introduction of corrections to the migration path,^19^ is an autonomous process that has also been observed in the absence of chemokine signaling when PGCs migrate in random directions.^18^ While the establishment of front-rear polarity in PGCs and other cell types has been extensively studied,^20–24^ it remains unclear how cell polarity is lost. Critically, the molecular network responsible for the periodic transition between run and tumble phases in eukaryotic cells is unknown.

Here, we identify the cellular and molecular cascade that leads to loss of cell polarity in PGCs, which contributes to accurate arrival at the migration target. Combining experiments and mathematical modeling, we determine that run and tumble behavior results from the periodic subsidence and reinstitution of a polar pattern, a behavior we term “polar pulsations”. Finally, we provide evidence that migrating PGCs lose their memory of cell polarity via non-directional blebbing that occurs during tumble phases, thus erasing pre-existing directional bias and allowing for changes in the migration path.

## Results

### Loss of front-rear polarity is initiated at the leading edge of the cell

To understand the intrinsic molecular mechanisms that control the transition from polar run into apolar tumble phases, we first investigated the dynamics of polarity loss in PGCs that migrate within embryos knocked down for the guidance cue Cxcl12a (Figure 1A).^11^ To this end, we labeled F-actin, a molecule enriched at the leading edge of migrating PGCs (Figure 1B, Lifeact, green, images 1, 3, and 5).^20^ Similar to previous reports,^18,20^ we found that, as judged by the polymerization of actin, the cells exhibit run (Figure 1B, images 1, 3, and 5) and tumble behavior (Figure 1B, images 2 and 4; Video S1) at around 12 min intervals. To characterize the transitions between run and tumble phases we measured the polarity of the actin signal over time (“Polarity Value”, see STAR Methods, Figure 1B, Graph; Video S1). We found that the depolymerization of actin at the cell front occurs before the morphological depolarization, suggesting that a reduction in actin polymerization is important for initiating polarity loss. In turn, consistent with the idea that Rac1 activity controls the pattern of actin polymerization,^20,25^ the dynamics of actin distribution mirrored that of activated Rac1 (Rac1 GTP, Figure 1C and 1D; Video S2), and the depolarization of activated Rac1 occurred before, or at the same time as, that of actin (Figure 1G).

**Figure 1.**
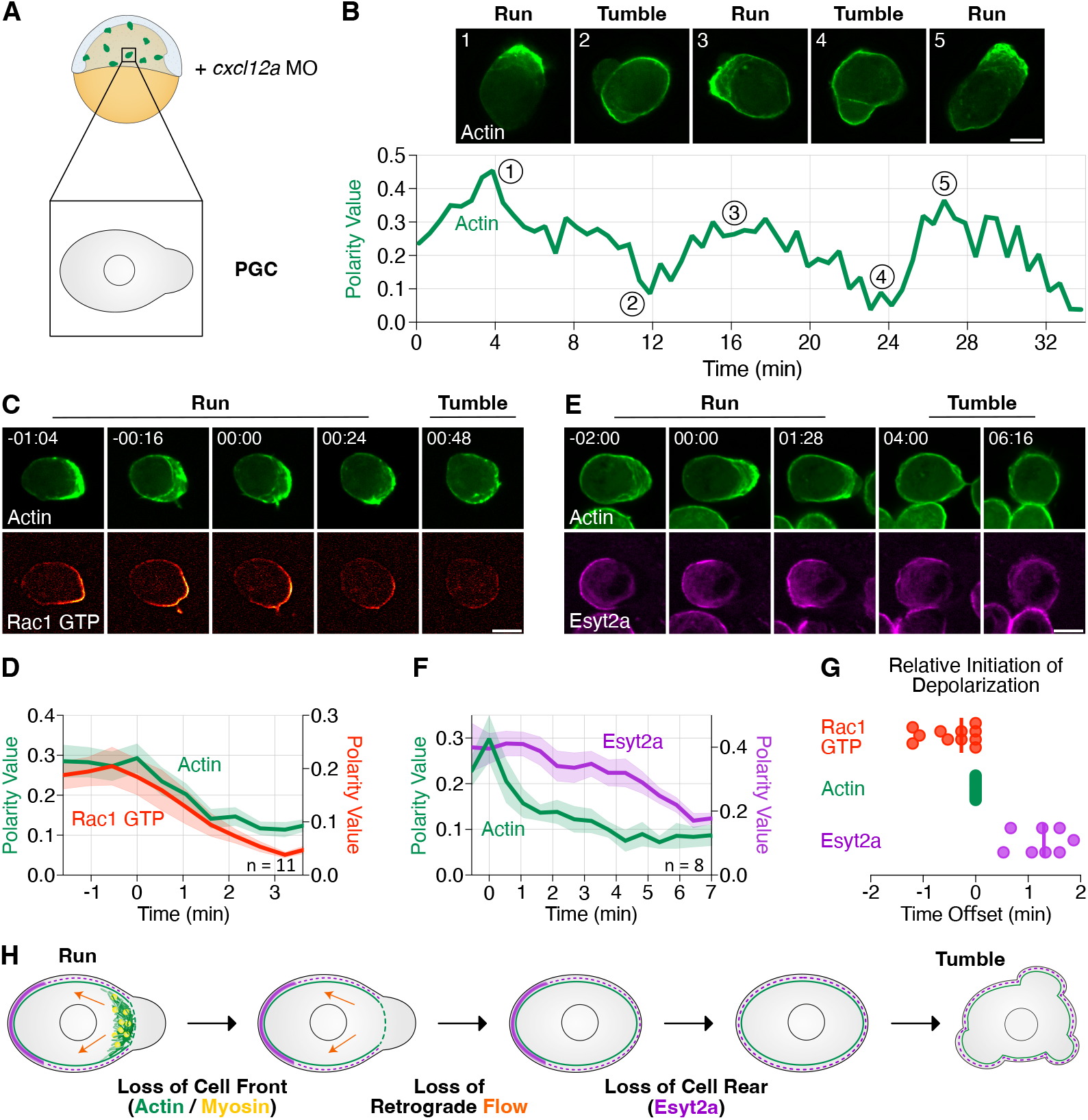
Distribution of molecules during run to tumble transitions. (A) Illustration of the experimental setup. Injection of a translation-blocking morpholino (MO) directed at the RNA encoding for the chemokine Cxcl12a results in non-directional migration of PGCs. (B) Dynamics of actin distribution during run and tumble phases (Lifeact, green, upper 5 panels) and the corresponding polarity value of actin over time (graph, with the 5 time points indicated, Video S1). (C) Images of a migrating PGC during run to tumble transition, presenting the distribution of actin (Lifeact, green) and GTP-bound Rac1 (Pak1 GBD, red). Time in mm:ss format (Video S2). (D) A plot showing the average polarity value of actin (green) and GTP-bound Rac1 (red) over time (see STAR Methods). Shaded areas show the standard error of the mean (SEM). (E) Distribution of actin (Lifeact, green) and Extended synaptotagmin (Esyt2a, magenta). Time in mm:ss format (Video S3). (F) A plot showing actin (green) and Esyt2a (magenta) average polarity values over time. Shaded areas represent SEM. (G) Initiation of depolarization of Rac1 GTP (red) and Esyt2a (magenta) relative to the time of actin depolarization (green). (H) Illustration of the events at the cell front and rear during run to tumble transition. Actin is labeled in green, myosin in yellow, Esyt2a in magenta. Orange arrows represent retrograde flow, and the black circles represent the nucleus. Scale bars: 10 µm. n indicates the number of cells.

To further characterize the transition between run and tumble, we examined the distribution of extended synaptotagmin-like 2a (Esyt2a, magenta, Figure 1E; Video S3), which is located at the cell rear.^20^ Measuring the polarity value of actin and Esyt2a over time revealed a steep drop in actin polarity between t = 0 min and 1.5 min (Figure 1F), while depolarization of Esyt2a started later and required more time to reach a steady state (Figure 1F; Video S3). Together, actin depolarization at the cell front preceded that of Esyt2a at the cell rear by 1.2 ± 0.6 min (Figure 1G).

Interestingly, this sequence of events is the reverse of what was described during polarity establishment in PGCs.^20^ Specifically, polarized activation of Rac1 initially directs the formation of the actin brushes to the cell front. Thereafter, actin retrograde flow translocates molecules such as Esyt2a and Ezrin to the cell rear, where they inhibit bleb formation, resulting in robust front-back polarity.^20^ Consistently, our findings show that the transition from run into tumble phases is correlated with a strong reduction in both Rac1 activity and actin polymerization at the cell front (Figure 1H). Under these conditions, retrograde flow is interrupted such that Esyt2a is not enriched at any specific position, allowing the formation of protrusions all around the cell perimeter (Figure 1H). Thus, while high levels of actin polymerization support the establishment and maintenance of cell polarity,^20^ a reduction in actin polymerization is correlated with the drop in cell polarity and transition into tumble phase.

### Contractility-deficient cells exhibit polar pulsations in actin polymerization

To identify the core molecular components required for the transitions between run and tumble states, we expressed a dominant-negative form of the Rho kinase 2 protein (DN ROCK)^26^ in PGCs (Figures 2A and S1A). Hereafter, we refer to such cells as contractility- deficient PGCs (cdPGCs, Figures 2A and S1A). This treatment reduced the ability of the cells to form blebs (Figure S1B), as well as the myosin-dependent retrograde flow, processes that are considered important for cell migration.^26,27^ Intriguingly, based on polarized actin assembly, cdPGCs still exhibited transitions between run and tumble phases (hereafter, cdRun and cdTumble phases, Figure 2B, Lifeact, green; Video S4), albeit with shorter run phases than in control cells (Figure S1C and S1D). In addition, since the definition of the rear requires retrograde actin flow,^20^ we found that in cdPGCs, the proteins localized to the back of migrating cells were uniformly distributed during cdRun phases (Ezrin, magenta, Figure 2C and 2D; Video S5).

**Figure 2.**
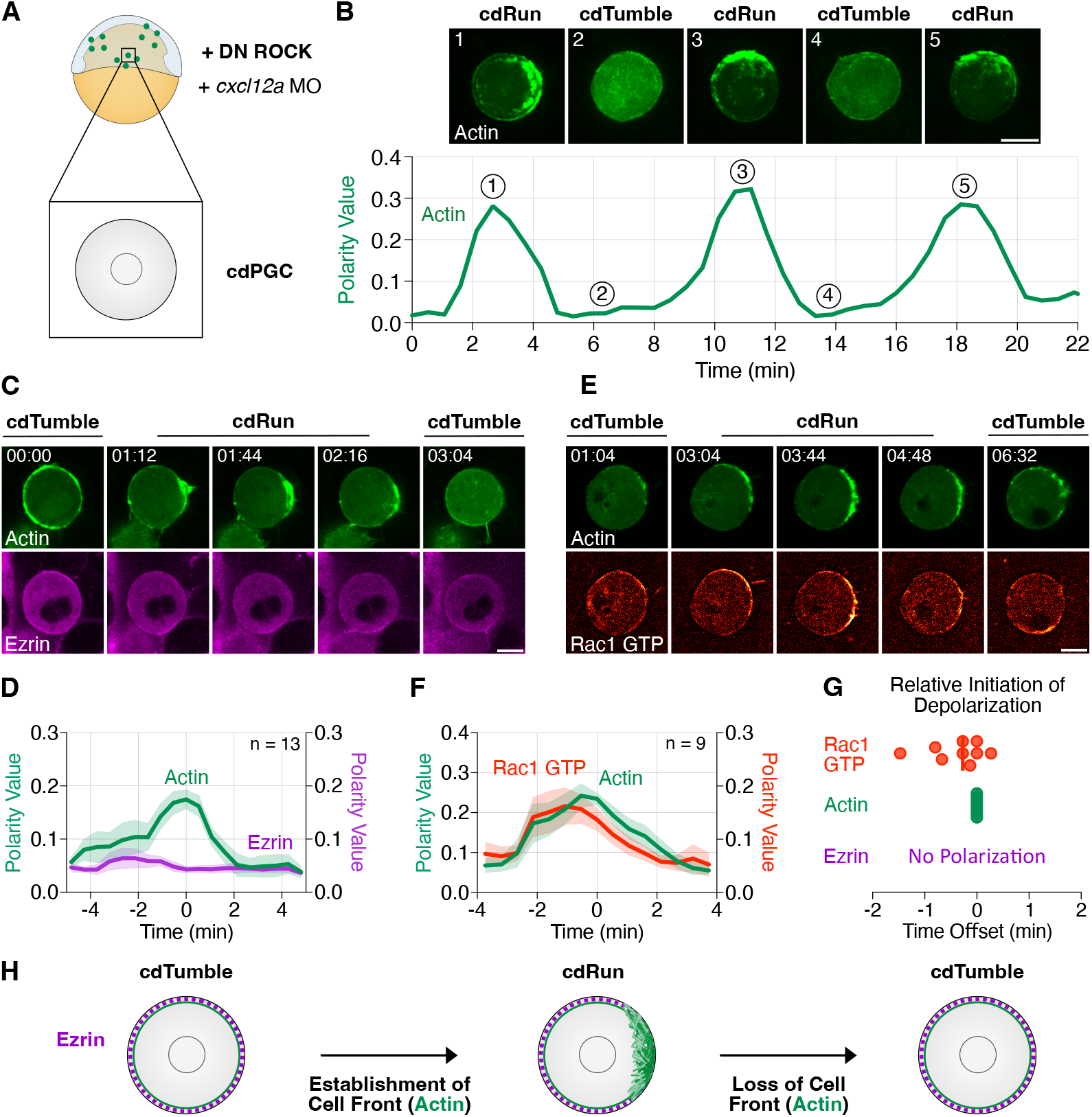
Actin polymerization oscillations in immotile contractility-deficient PGCs (cdPGCs) (A)Illustration of the experimental setup. Expression of a dominant-negative form of the ROCK protein (DN ROCK) and knocking down the expression of the chemokine Cxcl12a employing a morpholino antisense oligonucleotide (*cxcl12a* MO) generate contractility-deficient PGCs (cdPGCs) that are immotile and round. (B) Dynamics of actin distribution during run and tumble phases of cdPGC (Lifeact, green, upper 5 panels) and the corresponding polarity value of actin over time (graph, with the 5 time points indicated, Video S4). (C) Polarity oscillations in a cdPGC presenting actin (Lifeact, green) and Ezrin (magenta, Video S5). Time in mm:ss format. (D) A graph showing the average polarity value of actin (green) and Ezrin (magenta, Video S5) over time. Shaded areas represent SEM. (E) Distribution of actin (Lifeact, green) and Rac1 GTP (Pak1 GBD, red) in a cdPGC. Time in mm:ss format (Video S6). (F) A graph showing the average polarity values of actin (green) and Rac1 GTP (red) over time. Shaded areas represent SEM. (G) Initiation of depolarization of Rac1 GTP (red) relative to the time of actin depolarization (green). (H) A schematic presentation of polarity oscillations in cdPGCs. Actin is labeled in green and Ezrin in magenta. The black circle represents the nucleus. Scale bars: 10 µm. n indicates the number of cells.

These results show that the spatial distribution of polymerized actin correlates with run and tumble phases of PGCs independent of contractility, bleb formation, retrograde actin flow, motility, cell shape changes, and a defined cell rear, features that are associated with cell polarity establishment.^20,26,28–30^ The cdPGC thus allowed us to study the core mechanisms underlying the periodic gain and loss of cell polarity, a behavior we term “polar pulsations”.

### Rac1 controls actin polymerization in cdPGCs

To determine whether pulsations in polarized actin accumulation could be controlled by Rac1, we examined the spatial distribution of its activity in cdPGCs. Indeed, we observed polar pulsations of both actin and active Rac1 at the same location (Figure 2E), while the total Rac1 activity level was similar to that observed in control PGCs (Figure S2A). Specifically, the active Rac1 signal was homogeneously distributed during cdTumble (Figure 2E, time points 01:04 and 06:32) and enriched at the region of actin polymerization during cdRun phases (Figure 2E, time points 03:04 to 04:48; Video S6). Quantifications of the average polarity value of actin and Rac1 GTP confirmed these observations (Figure 2F). Consistent with the idea that Rac1 regulates the transition from run to tumble phases, depolarization of active Rac1 slightly preceded or mirrored the time of actin disassembly in cdPGCs (Figure 2G), similar to observations in control cells (Figure 1G). Overall, cdPGCs displayed polar pulsations of actin and Rac1 GTP, as observed in control PGCs, but without the accumulation of proteins such as Ezrin at the cell rear (Figure 2H).

To test whether Rac1 could indeed control polar pulsations in cdPGCs as in control cells, we manipulated its activity. Expression of a dominant-negative Rac1 mutant (Rac1^N17^)^31^ significantly reduced Rac1 activity in both control and cdPGCs (Figure S2B). Furthermore, the cells were unable to polymerize actin upon knockdown of Rac1 activity and instead displayed continuous tumble-like behavior (Figure S2C and S2D; Video S7).^20^ Consequently, the average polarity value of actin did not go above the threshold of >0.1 (see STAR Methods) within a 10- minute time frame for both control and cdPGCs (Figure S2C and S2D; Video S7). Conversely, hyperactivation of Rac1 signaling by expression of a constitutively active version of Rac1 (Rac1^V12^)^32^ resulted in the persistent apolar polymerization of actin around the cell perimeter, and no polar pulsations of actin were observed (Figure S2E and S2F; Video S8).

As such, these experiments show that Rac1 activity and actin polymerization play equivalent roles in regulating the polar pulsations observed in contractility-deficient and control PGCs. Therefore, we then focused on analyzing the control over Rac1 activity in cdPGCs.

### The GEF Prex1 controls actin pulsations by regulating Rac1 activity

Rho GTPases such as Rac1 act as molecular switches whose activity can shift between GTP bound (active) or GDP bound (inactive).^33^ The activity of Rho GTPases is regulated by proteins referred to as guanine nucleotide exchange factors (GEFs) and GTPase-activating proteins (GAPs).^34^ Based on deep sequencing of PGC transcriptome data and upstream signaling regulators in PGCs, we chose to examine the role of the GEF Prex1^35,36^ in PGC migration (Figure S3A). We tested the effect of two different dominant-negative forms of the protein in PGCs, namely the GEF activity-deficient mutant Prex1^E35A,N215A^ (hereafter referred to as DN Prex1)^37^ and Prex1 ΔDH (DN Prex1^ΔDH^), in which the GEF activity domain was deleted.^38^ The expression of either mutant protein significantly impaired PGC arrival at the developing gonad region (Figure S3A and S3B), indicating that Prex1 function is important for PGC migration. cdPGCs expressing DN Prex1 exhibited a reduction in actin polymerization and displayed persistent tumble-like behavior, as indicated by an average polarity value of actin <0.1 (Figure 3A; Video S9). While most of the control cells polarized at least once within a 10 min time frame (60.0 %), only a small fraction of DN Prex1-expressing cdPGCs did so (17 %, Figure 3C). Consistent with the role of Prex1 as a GEF, Rac1 activity in cdPGCs was reduced upon knockdown of Prex1 function (Figure 3B). Interestingly, overexpression of wild-type Prex1 led to an increase in the amount and persistence of actin polymerization (Figure 3D; Video S10), and Rac1 activity (Figure 3E), resulting in longer run phases (Figure 3D and 3F; Video S10). This effect was even more pronounced upon expression of a constitutively active version of Prex1 (CA Prex1, Figure 3G; Video S11). We observed persistent apolar polymerization of actin in multiple cdPGCs and a marked increase in Rac1 activity upon expression of CA Prex1 (Figure 3H). Overall, our experiments suggest that Prex1 positively regulates Rac1 activity and actin polymerization and, thus, could regulate the polar pulsations in cdPGCs.

**Figure 3.**
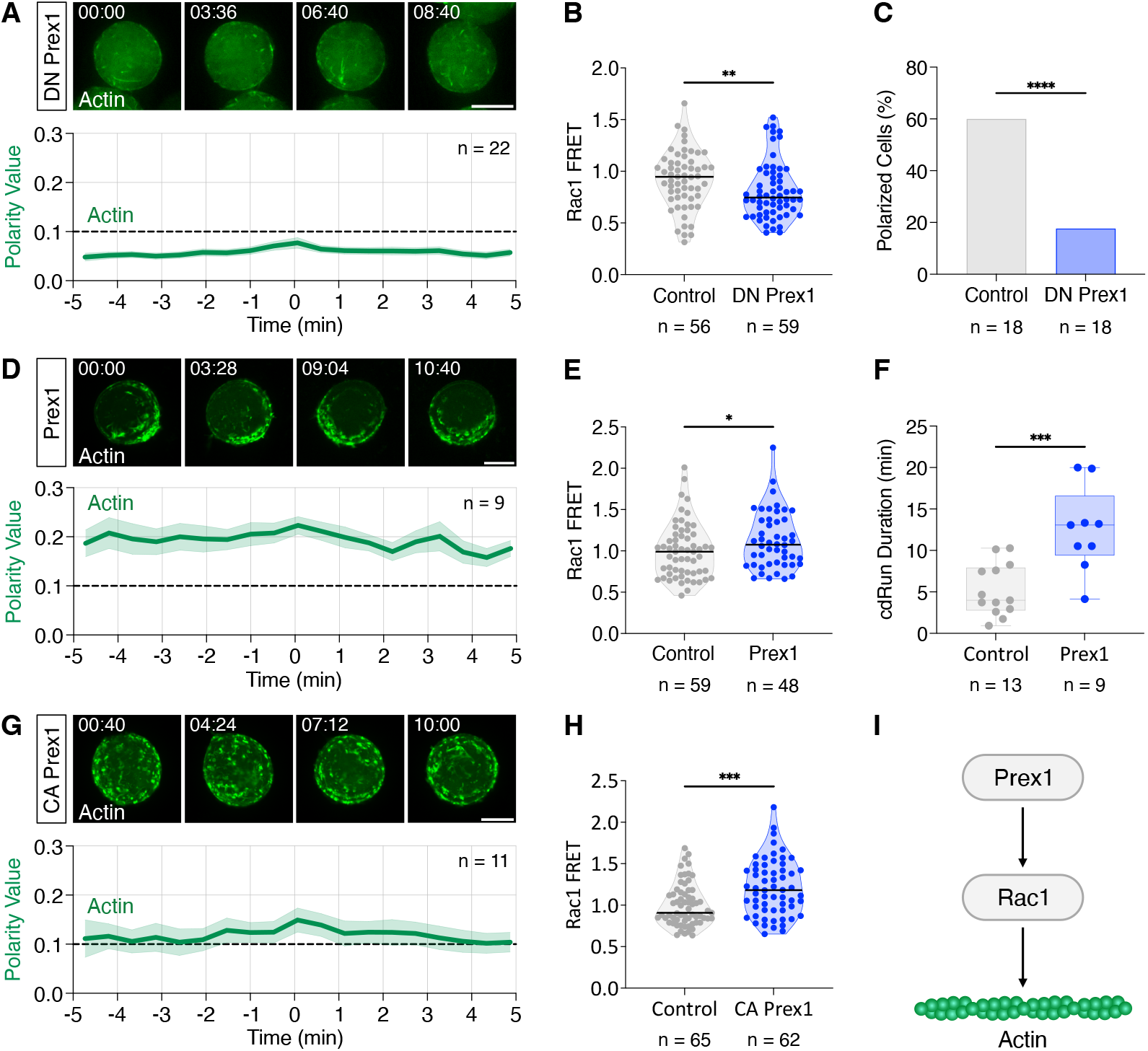
The GEF Prex1 controls Rac1 activity and actin polymerization in cdPGCs. (A)Dynamics of actin distribution (Lifeact, green) in a cdPGC expressing the dominant- negative form of Prex1 (DN Prex1, Video S9). The graph presents the average polarity value of actin over time (see STAR Methods). Shaded areas represent SEM. (B) Normalized Rac1 FRET value for control cdPGCs and in cdPGCs expressing DN Prex1. **, P- value = 0.083 (Mann-Whitney test). (C) Fraction of polarized control cdPGCs and DN Prex1-expressing cdPGCs. ****, P-value < 0.0001 (Chi-square test). (D) Dynamics of actin distribution (Lifeact, green) in a cdPGC overexpressing Prex1 (Video S10). The graph shows the average polarity value of actin over time. Shaded areas represent SEM. (E) Rac1 activity level (calculated by FRET ratio) in control cdPGCs and in cdPGCs overexpressing Prex1. *, P-value = 0.0403 (Mann-Whitney test). (F) Duration of run phases for control cdPGCs and cdPGCs overexpressing Prex1. ***, P-value = 0.0003 (Mann-Whitney test). (G) Dynamics of actin distribution (Lifeact, green) in a cdPGC expressing constitutively active Prex1 (CA Prex1, Video S11). The graph shows the average polarity value of actin over time. Shaded areas represent SEM. (H) Rac1 activity level (calculated by FRET ratio) in control cdPGCs and in cdPGCs expressing CA Prex1. ***, P-value = 0.0003 (Mann-Whitney test). (I) A schematic representation of the signaling cascade controlling actin polymerization in cdPGCs. F-actin is presented as chains of green spheres. Scale bars: 10 µm. The dashed lines in panels A, D, and F represent the threshold set to define apolar tumble phases (<0.1) and polar run phases (>0.1). Time in mm:ss format. n is the number of cells examined.

### The Pak1 protein controls the frequency of actin polar pulsations

As presented above, actin polymerization is controlled by the proteins Prex1 and Rac1 (Figure 3I). However, such a linear signaling cascade would not account for the polarity pulsations we observe. Rather, an oscillatory or pulsatory behavior requires a negative feedback loop, which brings the signaling network back to its starting point,^39^ coupled with non-linearity in the reactions to prevent the system from reaching a steady state.^40,41^ Indeed, the GEF activity of Prex1, which depends on its localization to the plasma membrane, can be regulated by post-translational modifications such as phosphorylation. Relevant to this work, cell culture experiments have shown that p21-activated kinase (Pak1) can inhibit the activity of Prex1 and Prex2.^42,43^ In addition, Pak1 can be activated by Rac1,^44^ thereby establishing a negative feedback loop (Figure 4A) that was suggested to modulate signaling levels downstream of plasma membrane receptors.^42,43^ Accordingly, we next studied the role of Pak1 in the context of polar pulsations of actin accumulation (Figure 4A–A’’’).

**Figure 4.**
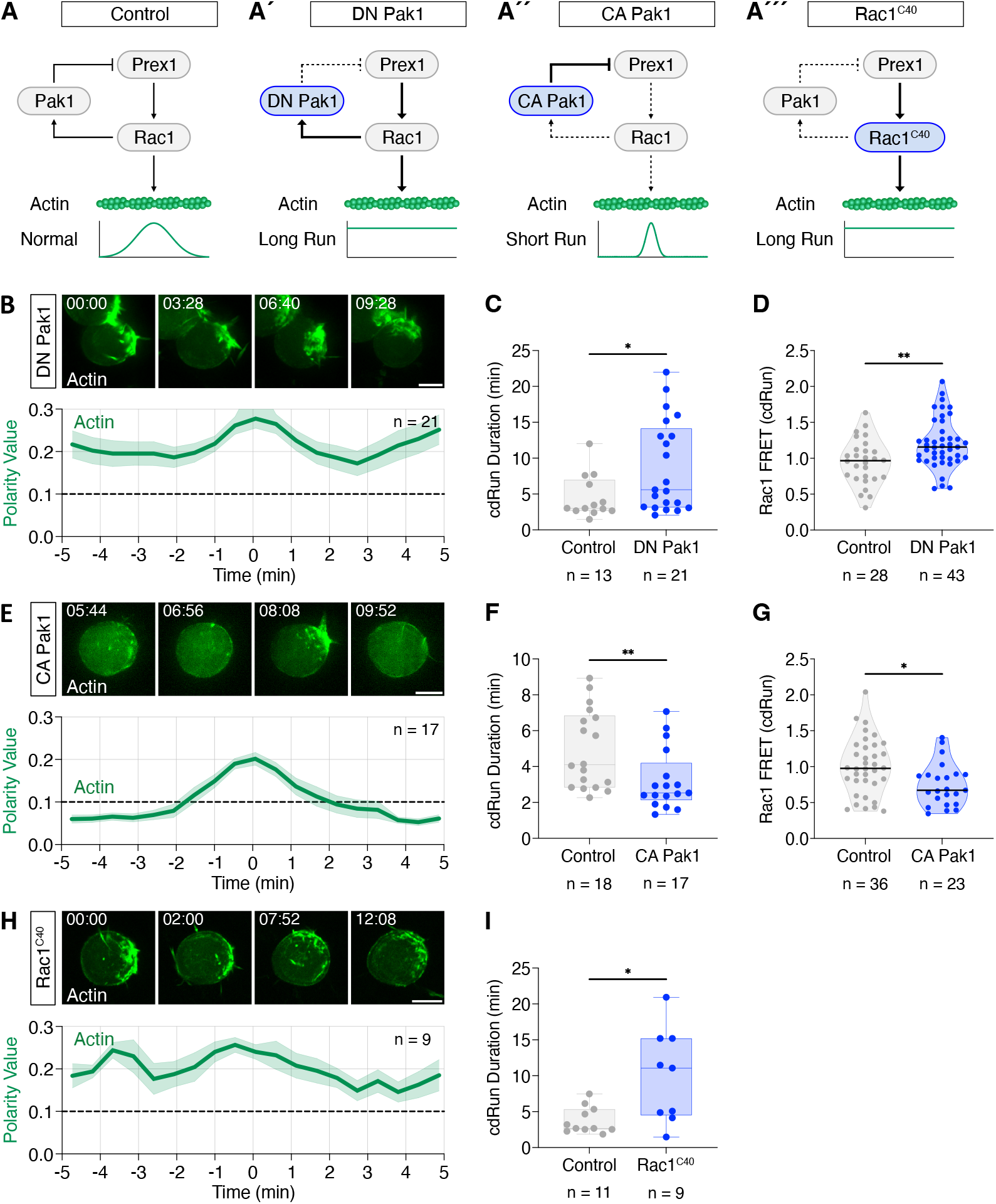
A Pak1-mediated negative feedback loop regulates polarity oscillations in cdPGCs. (A–A’’’) Schematic representations of the negative feedback loop in cdPGCs, with the manipulated molecular components highlighted in blue, and graphs presenting the expected effects of the treatments. Pointed arrows (→) represent activation, and blunt arrows (− I) indicate inhibition. Dashed arrows represent a decrease in function, and bold arrows an increase. F-actin is presented as chains of green spheres. (B) Dynamics of actin distribution (Lifeact, green) in a cdPGC expressing the dominant- negative form of Pak1 (DN Pak1, Video S12) with the graph showing the average polarity value of actin over time (see STAR Methods). Shaded areas represent SEM. (C) Duration of run phases for control cdPGCs and for cdPGCs expressing DN Pak1. *, P-value = 0.0298 (Mann-Whitney test) (D) Rac1 FRET value during cdRun phases for control cdPGCs and in cdPGCs expressing DN Pak1. **, P-value = 0.0029 (Mann-Whitney test). (E) Dynamics of actin distribution (Lifeact, green) in a cdPGC expressing a constitutively active form of Pak1 (CA Pak1, Video S13), with the graph below showing the average polarity value of actin over time. Shaded areas represent SEM. (F) Duration of run phases for control cdPGCs and cdPGCs expressing CA Pak1. **, P-value = 0.0077 (Mann-Whitney test). (G) Rac1 FRET values during cdRun phases for control cdPGCs and for cdPGCs expressing CA Prex1. *, P-value = 0.0124 (Mann-Whitney test). (H) Dynamics of actin distribution (Lifeact, green) in a cdPGC expressing Rac1^C40^ (Video S14), with the graph below showing the average polarity value of actin. Shaded areas represent SEM. (I) Duration of run phases for control cdPGCs and for cdPGCs expressing Rac1^C40^. *, P-value = 0.0124 (Mann-Whitney test). Scale bars: 10 µm. The dashed lines represent the threshold set between apolar tumble phases (<0.1) and polar run phases (>0.1). Time in mm:ss format. n represents the number of cells.

Consistent with the proposed negative feedback loop, cdPGCs expressing the dominant-negative Pak1^K331R^ mutant (DN Pak1)^45^ displayed more persistent actin polymerization (Figure 4A’ and 4B; Video S12). Accordingly, in contrast to control cdPGCs (see Figure 2B), cells expressing DN Pak1 had longer cdRun phase durations (Figure 4C) and showed elevated Rac1 activity during these phases (Figure 4D). Yet, Pak1 knockdown did not affect the average cdTumble phase duration nor Rac1 activity during cdTumble phases (Figure S4A and S4B). Furthermore, cdPGCs that express a constitutively active version of Pak1 (CA Pak1, Pak1^T455E^)^46^ showed very short cdRun phases (Figure 4A’’, 4E, and 4F; Video S13) and a corresponding decrease in Rac1 activity (Figure 4G). Interestingly, hyperactivation of Pak1 also significantly increased the duration of cdTumble phases due to a reduction of Rac1 activity (Figure S4A and S4B). Similar results were obtained when employing another version of CA Pak1, namely Pak1^T455E,H82,85L^ (hereafter CA Pak1^H82,85L^, Figure S4C), which is unable to bind Rac1 and be regulated by it,^46^ indicating that Pak1 controls the duration of run phases independent of potentially sequestering Rac1 in cdPGCs. These results are consistent with the idea that Pak1 function during run phases is mediated by its kinase activity, which regulates Rac1 activity by inhibiting Prex1 (Figure 4A).

To further examine the negative feedback loop downstream of Rac1, we expressed in cdPGCs the Rac1^C40^ mutant that cannot bind and activate Pak1.^47^ This manipulation leads to a situation in which Prex1 activates Rac1^C40^ without activating the Pak1-mediated negative feedback loop (Figure 4A’’’). Indeed, in this experimental setup, the duration of cdRun phases was increased (Figure 4H and 4I; Video S14), similar to the findings in Pak1 knockdown cells (Figure 4B and 4C). The Rac1^C40^-induced phenotype was not a result of increased Rac1 protein amount or activity *per se*, since overexpression of wild-type Rac1 did not increase the duration of cdRun phases in cdPGCs (Figure S5A and S5B; Video S15). Thus, the data reveal evidence for a negative feedback loop in which the kinase Pak1 regulates the duration of run phases and, thus, the frequency of actin polarity pulsations downstream of Rac1, presumably by inhibiting Prex1 activity (Figure 4A).

### Modifying the polar pulsation parameters impairs PGC arrival at the gonad region

We found that manipulation of Pak1 signaling alters the frequency of polarity pulsations in cdPGCs. Notably, the frequency of run and tumble transitions is important for the arrival of germ cells at the gonad by allowing them to correct their migration path, especially when they are close to their target.^18,19^ To examine whether the molecular network presented in Figure 4A also operates in cells whose contractility has not been modified, we manipulated the function of Pak1 proteins in otherwise normal PGCs. Indeed, the expression of DN Pak1 or CA Pak1 increased the number of ectopic cells found outside the gonad region at 24 hours post fertilization (hpf) (Figure 5A and 5B). Thus, pulsations at frequencies that are too low (DN Pak1) or too high (CA Pak1) impair the ability of PGCs to reach their target.

**Figure 5.**
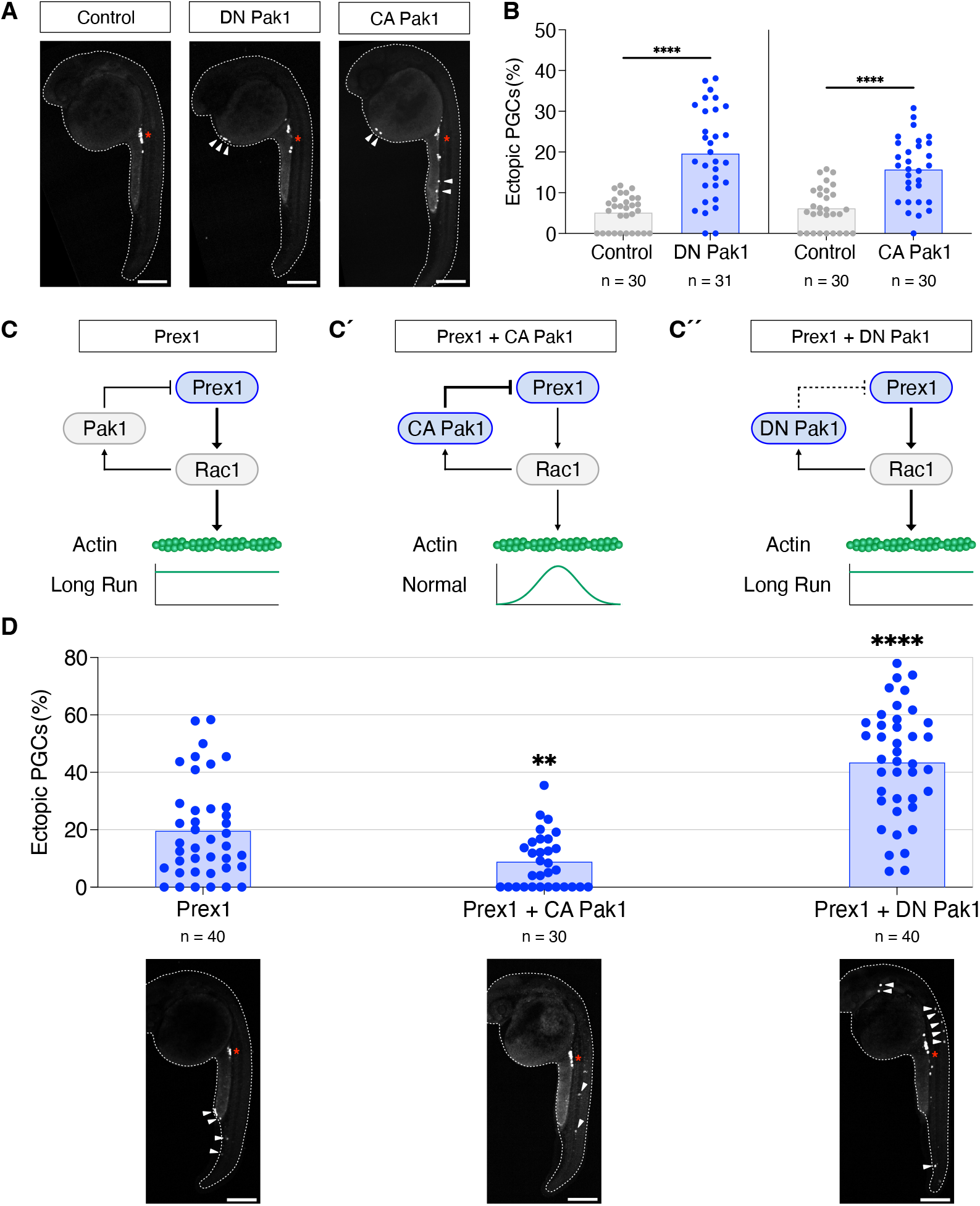
Manipulation of polarity oscillations impairs PGC arrival at the gonad region. (A)24 hpf embryos with GFP-labeled PGCs that express either control, DN Pak1 or CA Pak1 protein. The migration target sites are marked by red asterisks and the white arrowheads point at ectopic cells. (B) The fraction of ectopic PGCs induced by expression of either control, DN Pak1 and CA Pak1 proteins. ****, P-value < 0.0001 (Mann-Whitney test). **(C–C’’)** Schematic representations of the negative feedback loop that functions in PGCs. The manipulated molecular components are highlighted in blue, and the graphs present the expected outcome of the manipulations. Pointed arrows (→) represent activation, while blunt arrows (?I) indicate inhibition of protein function. Dashed arrows indicate reduced function, and bold arrows an increase. F-actin is labeled as chains of green spheres. **(D)** The fraction of ectopic PGCs in 24 hpf embryos, with representative embryos shown below each column. The migration target sites are labelled with asterisks, and the arrowheads point at ectopic GFP-labeled PGCs. **, P-value = 0.0037; ****, P-value < 0.0001 (Mann-Whitney test). Scale bars: 200 µm. The white dashed lines indicate the contours of the embryos. n is the number of cells examined.

As a further test for the proposed signaling loop and the potential of its components to control the frequency of polarity pulsations and arrival at the target, we designed experiments in which the activity of more than one molecule was affected. As presented above, overexpression of Prex1 increased the duration of run phases (Figures 3D, 3F and 5C). According to the proposed signaling network, enhancing the negative feedback by expressing CA Pak1 should revert this phenotype (Figure 5C’), while cells that are concurrently knocked down for Pak1 should display an even more severe phenotype (Figure 5C’’). To achieve a stronger effect of Prex1 overexpression, we co-expressed G protein subunits beta 1 and gamma 2 (G*β*1γ2), a known upstream activator of Prex1.^37^ As expected, adding CA Pak1 significantly decreased the level of mismigration observed in PGCs expressing the activated Prex1 protein (Figure 5D). The proposed cascade is also supported by the finding that expressing activated Prex1 and DN Pak1 led to a very high level of ectopic PGCs compared to the phenotype observed when expressing each one alone (Figure 5B and 5D).

Together, these experiments suggest that Pak1 and Prex1 function in the same signaling pathway and that Pak1 inhibits Prex1 activity in zebrafish PGCs (Figure 4A). Different levels of the ligand that affect G*β*γ signaling and, thus, Prex1 activity could potentially alter the balance between positive and negative signals. In this way, receptor activation could control the frequency of polar pulsations to ensure proper arrival at the target.

### Circuit theory of polar pulsations

To gain mechanistic insight into the dynamical processes that control polar pulsations, we developed a minimal mathematical model of the core biochemical circuit (Figure 6A). The dynamics can be captured by three fields, 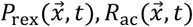 and 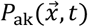,which represent respectively the local concentrations (number density) at position 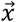 and time *t* of the activated proteins of Prex1, Rac1 and Pak1 along the membrane. For simplicity, we assumed that the inactive form of the proteins is kept at a roughly constant cytosolic concentration. Moreover, since the concentration of active Rac1 is strongly coupled to Prex1 in the limit of interest, up to a constant scaling factor it can be replaced by the active Prex1 concentration. Then, when the concentrations of active proteins is high, stochastic number fluctuations can be neglected and the biochemical reactions can be captured by a set of rate equations, which take the form of a coupled reaction-diffusion system (see STAR Methods),

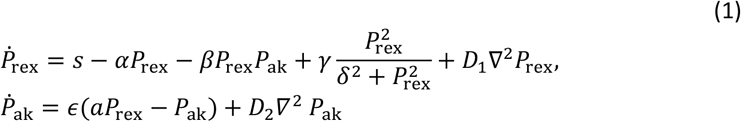

**Figure 6.**
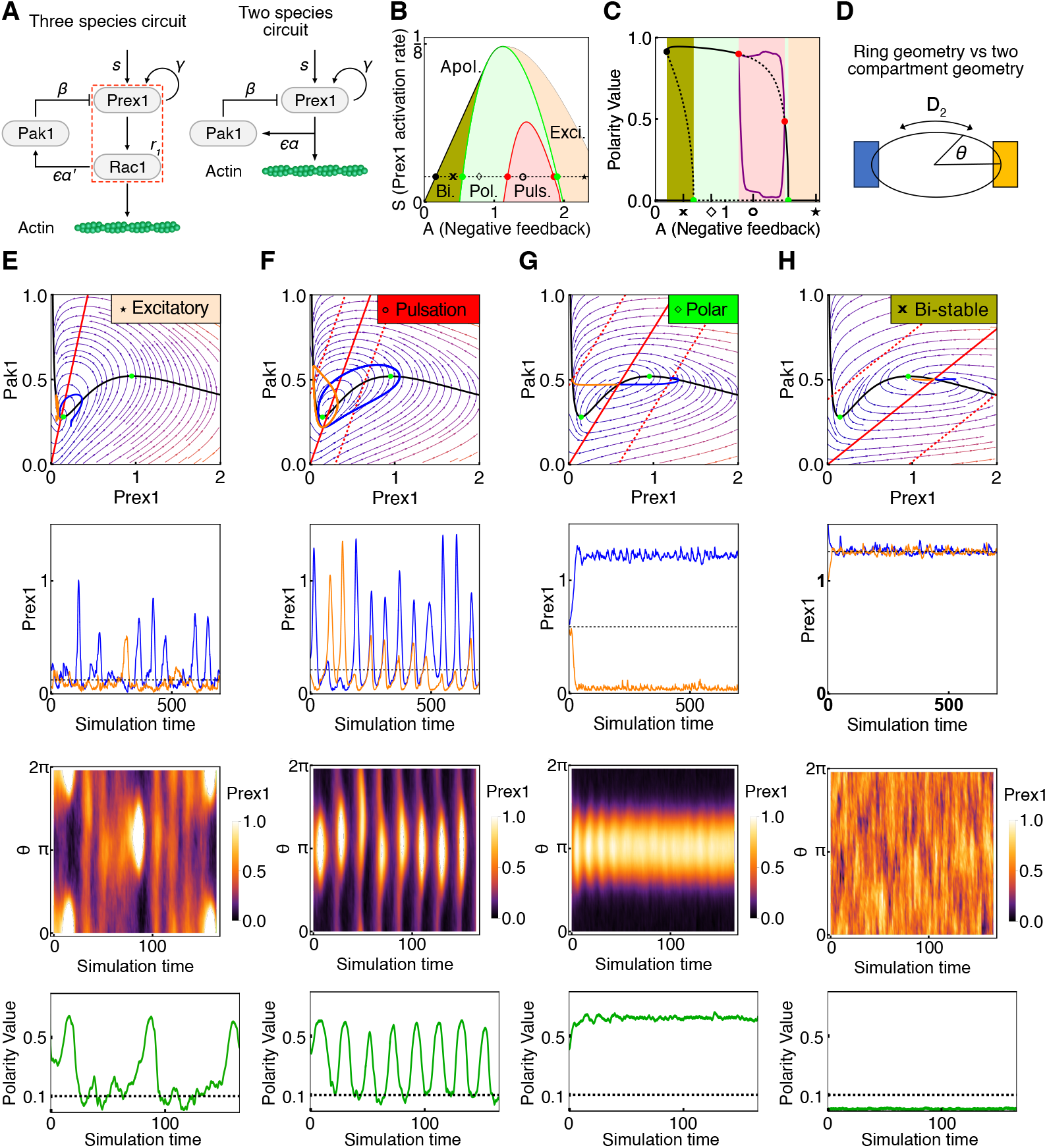
Phase behavior of the mathematical model for the biochemical loop. (A)Illustration of the three species biochemical network, its reduced two species counterpart, and the corresponding terms in Eq.(1). (B) Phase diagram in the S − A plane for the two-compartment model (7) at ϵ = 0.067 and D_2_ = 5. The black green and red lines denote Saddle node, Pitchfork, and Hopf bifurcations, respectively. (C) Bifurcation diagram along a constant S = 0.02 cross section of the phase diagram (denoted by thin dashed line in panel (B)) in terms of the polarity value between the two compartments 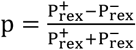. The black solid lines are stable steady states. The purple upper (lower) curves denote the maximal (minimal) polar amplitude within the polar pulsations phase. Dashed lines are unstable steady states. (D–G) top row: phase portrait for the flow prescribed by the reaction terms in (3) in the excitatory, pulsation, polar and stable apolar states of the two compartment model (7). The corresponding values for A for each plot are marked by the symbols in (B) and (C). Black and red curves are the Prex1 and Pak1 nullclines (4) respectively. Dashed red lines are the shifted Pak1 nullclines (9) of polar states. The green dots denote pitchfork bifurcations as in (B) and (C). Orange and blue curves in the top and second row are numerical solutions for the time varying local compositions in the + and – compartments. Third and last rows present numerical solutions of the continuous space ring geometry diagram along the same constant S = 0.02 cross section of the phase diagram, and similar values of A as for the two compartment model but with at ϵ = 0.07, D_1_ = 0.001 and D_2_ = 0.1. The polarity value in the bottom row was computed as in the experiment. Simulations were supplemented with a Gaussian white noise ξ for the activation term S → S + σ^2^ξ with amplitude σ = 0.15 for the two compartment model and σ = 0.2 for the continuous space mode (where the noise was Gaussian and white in both space and time).

Equations (1) belong to the FitzHugh–Nagumo class of dynamics,^48,49^ with Prex1 and Pak1 playing the role of excitatory and inhibitory species, respectively. Here, Prex1 is locally activated by upstream membrane-bound regulators such as G*β*γ proteins^35^ that account for the response to external chemical gradients, if present (Figure 6A). In their absence, we assume a uniform activation rate of Prex1 along the membrane that, away from some limiting activation capacity, enters the Prex1 dynamics as a positive rate constant *s*. Saturation at maximal capacity is absorbed into the spontaneous deactivation term −*αP*_ref_. The inter- species reaction terms are based on our knowledge of the corresponding reactions (see STAR Methods). Active Prex1 is locally deactivated by active Pak1, as captured by the nonlinear term −*βP*_ref_*P*_ak_, while Pak1 is activated by active Rac1, as captured by the term ϵ*aP*_ref_. (Note that, here, in the limit of interest, the concentration of Rac1 has been replaced by Prex1.) Moreover, in common with Prex1, Pak1 is subject to spontaneous deactivation, as captured by the term −ϵ*P*_ak_. Lastly, the observed pulsating dynamics indicate that the biochemical loop presented in Figure 4A also includes a positive feedback loop, which we model as a Hill-type function for the Prex1 dynamics. While this choice is motivated by previous work,^50–52^ within some mild requirements, the precise functional form of such a positive feedback loop does not impact significantly on the behavior of the system (see STAR Methods). Finally, the dispersion of all components is taken to be diffusive with respective diffusion coefficients *D*_1,2_.

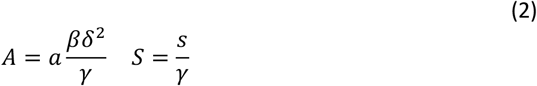

The model supports different dynamical regimes that are summarized in Figure 6B and 6C, and displayed in Figure 6E–H for two geometries: a simplified yet fully tractable model comprising only two spatial compartments (top two rows); and a more realistic continuous ring geometry (bottom two rows). Both show the same qualitative behavior. The transitions between these dynamical regimes are fully determined by two dimensionless control parameters which encode the strength of negative feedback on Prex1 and the rescaled Prex1 activation rate, respectively (Figure 6B). With Pak1 characterized by much faster diffusion *D*_2_ ≫ *D*_1_^53,54^ (as active Prex1 is membrane-bound^55^), yet delayed response^43,56^ (small ϵ), the system supports a dynamical regime of localized excitations (Figure 6E). Indeed, this hallmark of FitzHugh–Nagumo dynamics has been a focus of prior studies modeling polarity in other types of chemotactic cells.^57–60^ Here, a quiescent low Prex1-activity apolar state is stable (cdTumble); but when noise-driven above a threshold, it undergoes polar excursions (cdRun phases) with broadly distributed waiting times in between.^59,61^ This excitatory regime only appears in our model at strong enough negative feedback *A* (Figure 6B), and is realized experimentally in cdPGCs expressing CA Pak1 (Figure S6, see also STAR Methods). Upon decreasing *A*, inhomogeneous polarized states become stabilized. Here, the fast dispersion of Pak1 from regions of high Prex1 activity swamps the system and suppresses regions of low activity, leading to a stable polarized pattern. This phenomenon represents a Turing pattern, and it emerges here due to the large disparity between the diffusivity of the species, a situation not considered in previous studies (see STAR Methods).

A further reduction in *A* leads to a regime marked by temporal pulsations of the polar pattern (Figure 6F), a phase also not encountered in previous studies. Note that at low ϵ, the excitatory regime transitions directly into polar pulsations (where the narrow green region at large *A* vanishes, Figure 6B and 6C). This phase can be described as the interference of the aforementioned Turing instability, which stabilizes stable polar patterns, with a limit cycle instability (also known as a Hopf bifurcation). The latter is a Hallmark of the spatial-less FitzHugh–Nagumo model, whereupon tuning of parameters can lead to periodic oscillations in activity. The coincidence of the two instabilities gives rise to the observed periodic pulsations of a polar pattern.

Indeed, we find that this regime fully captures the phenomenology of cdPGC, while the distinct phenotypes that have been observed upon protein activity manipulation, fall naturally within the other dynamical regimes predicted by the model.

Upon further decreasing *A*, or increasing the activation rate *S*, the polar pulsations are abolished, and a stable polar state emerges (Figure 6G). Indeed, the three treatments of DN Pak1, overexpressing Prex1, and Rac1^C40^ shift the system toward this regime, where the experimental behaviors are marked by persistent polar states (long cdRun phases) occasionally interrupted by noisy apolar transients (cdTumble phases, Figure S7, see also STAR Methods).

When *A* is further decreased or *S* increased, an apolar state becomes stable (Figure 6H), coexisting with the stable polar state (Figure 6B and 6C, bistable region). Finally, upon a further decrease in *A*, only the apolar state remains stable (Figure 6B and 6C). Indeed, the disappearance of a stable polar pattern, termed a saddle-node bifurcation, was at the focus of a recent line of theoretical work that proposed a novel mechanism for adaptive memory in chemotactic cells.^62^

### Blebbing during tumble phases is important for erasing polarity axis *memory*

In the context of a response to a stimulus, cellular “memory” refers to the ability to maintain certain features, such as front-rear polarity, for an extended time, even after the initial signal is no longer present.^63,64^ In the absence of guidance cues, migratory PGCs are found to initiate run phases in random directions following tumble phases.^18^ This indicates that the polarity and its memory are erased during tumble phases,^18^ rendering the cell naïve and able to introduce corrections to its migration path.

To investigate the phenomenon of cellular memory loss, we examined the potential role of blebs during tumble phases in the process. First, we quantified the angle between the polarity vectors of two consecutive run phases, as measured by the polarity value of actin (see STAR Methods). Similar to our previous findings,^18^ we found that in the absence of guidance cues after tumble phases, migratory PGCs formed a new cell front at an average angle of around 80°, which reflects a small deviation from a completely random distribution (Figure 7A and 7C). In contrast, in cdPGCs, the formation of the new cell front occurred close to the previous one, at an average angle of around 50°, indicating a significant bias (Figure 7B and 7C). Indeed, this finding is supported by the mathematical model (Figure 6E).

**Figure 7.**
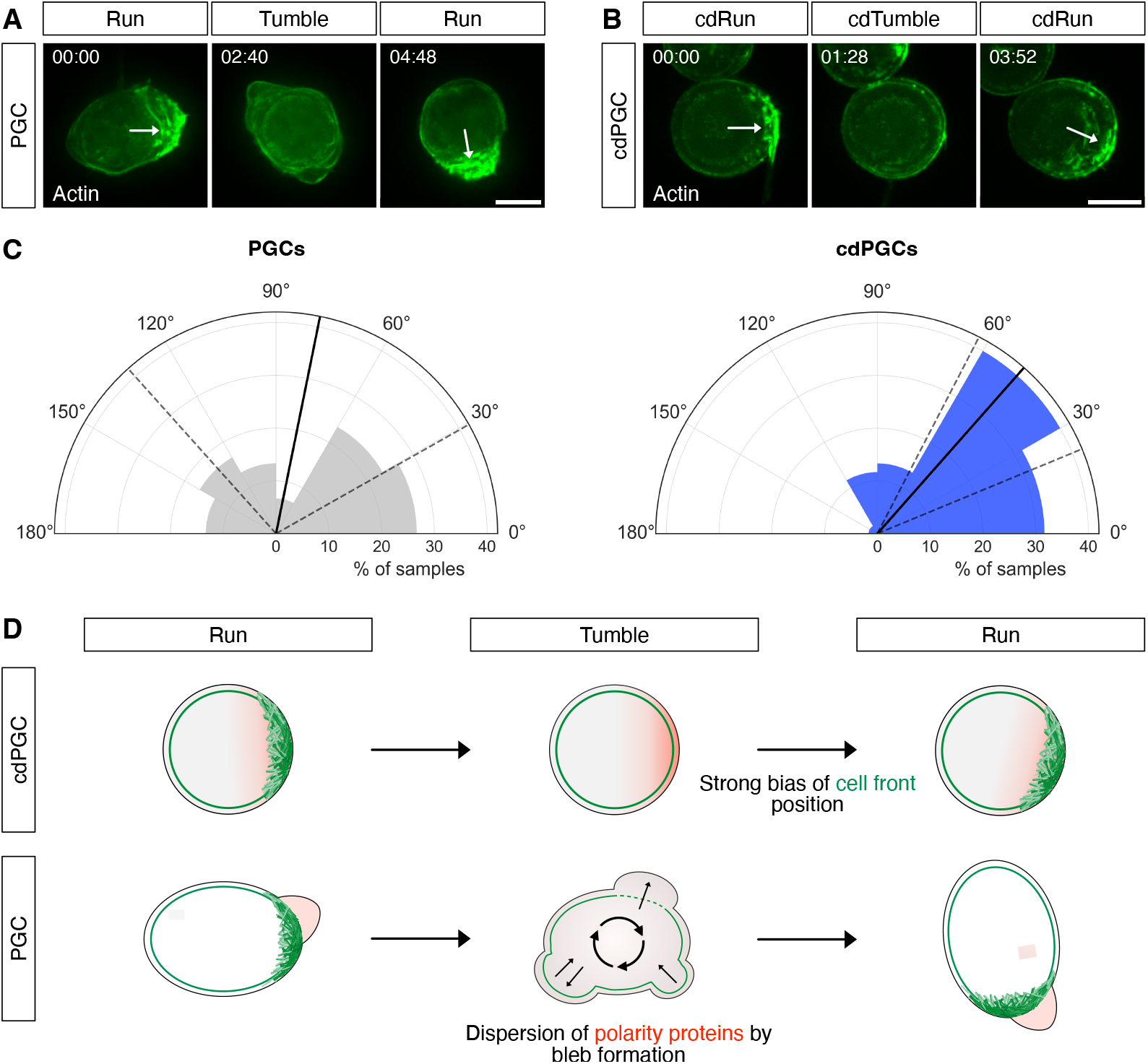
Bleb formation during tumble phases erases molecular memory in PGCs. (A, B) The direction of the polarity vector for consecutive run phases interspaced by a tumble phase of a representative (A) control PGC and (B) a contractility-deficient PGC (cdPGC, Lifeact, green). The white arrows indicate the direction of polarization vectors (see STAR Methods). Time in mm:ss format. Scale bars: 10 µm. (C) Rose plots showing the relative distribution of the angles between the vectors of the direction of polarization of consecutive run phases of PGCs or cdPGCs (white arrows in a and b, see STAR Methods). *, P-value = 0.044 (Mann-Whitney test). n = 30 (PGCs), n = 60 (cdPGCs), n is the number of cells. (D) An illustration showing the effect of blebbing on the preservation of migration-direction memory PGCs. In control PGCs, the polarity-promoting proteins, which regulate the formation and maintenance of the cell front, are dispersed during tumble phases by blebs, resulting in the formation of the new cell front in random or weakly biased directions. In contrast, the polarity-promoting proteins are not mixed in cdPGCs during cdTumble phases, leading to the formation of the new cell front in a direction that is biased towards the position of the preceding front. Actin is labeled in green, arrows illustrate the mixing of the cytoplasm, and red represents the distribution of polarity promoting factors.

These results revealed a putative cellular mechanism that erases the memory of front-rear polarity. While maintaining a consistent axis is important for persistent migration during run phases, polarity must be abolished to render the cell more responsive to guidance cues and modify its migration direction. We suggest that in addition to diffusion within the cell, the non-directional blebbing during tumble phases mixes those molecules that provide molecular memory, contributing to its periodic erasure (Figure 7D).

## Discussion

Periodic dynamics have been found in diverse cell types ranging from microbes^65,66^ to plants^67,68^ and mammalian cells^69^ and play central roles in controlling diverse cellular processes including the circadian rhythm,^70,71^ gene expression, differentiation, and mitosis.^72,73^ In the context of cell polarity and migration, periodic dynamics have been observed in the activity of Rac1,^74,75^ RhoA,^76^ and Cdc42,^77,78^ and regulate migration speed and arrival at the target.^79,80^

Here, we used zebrafish primordial germ cells as an *in vivo* model to study the periodic transitions between polar (migrating) and apolar (non-migrating) phases. Our results showed that run and tumble behavior results from polar pulsations, the periodic erasure and establishment of spatially patterned molecular activity. These pulses exhibit periodicity and are sustained over time; they are polarized to a single region of the cell and are stationary in nature, but can occur at random positions in the absence of external directional cues. Therefore, the polar pulsations are distinct from previously described mechanisms, such as apolar oscillations, which exhibit no spatial patterning;^81^ polar-reversal oscillations, that result in a periodic switch of polarity direction between two poles, reported in migrating cells;^66,75,82^ or propagating waves of signaling molecules in *Dictyostelium*.^83,84^ The polar pulsations in cdPGCs share similarities with oscillations of Cdc42 during bud formation in *Saccharomyces cerevisiae*.^77^ However, the Cdc42 oscillations are dampened and eventually establish a fixed polar pattern, which differs from the polar pulsations mechanism that operates in cells that maintain directed migration behavior. Notably, we show that the range of dynamic behaviors observed experimentally can be captured by a minimal mathematical framework that captures the core biochemical regulatory components. Although this model shares similarities with previous models of cellular polarity, the range of experimental parameters encountered here drives the system into regimes where new dynamic phases, including polar pulsations, appear.

We observed that pulsations in polarized Rac1 activity control periodic polymerization and depolymerization of actin and, consequently, the establishment and elimination of the cell front (Figure S8A and S8B, see also Figures 1H and 2H). To identify the relevant molecules and examine functional relationships among them, we first defined the components that are sufficient for the polar pulsations and those that are not essential. Intriguingly, many of the molecules and processes that are required for migration or correlated with it (e.g., myosin activity and actin retrograde flow) are not needed. We showed that changes in cell shape, blebbing, localization of proteins at the cell rear, active migration, and flow of the actomyosin cortex are not required for polar pulsations (Figure 2B and 2H). Instead, we identified the GEF Prex1, the kinase PAK1 and the Rho GTPase Rac1 that function in a negative feedback loop that controls the pulsations in polarized Rac1 activity and actin distribution (Figure S8A).

Intriguingly, the interactions among these molecules can be modulated to alter the pulsations frequency. For example, increasing the activity of Prex1 or Pak1 reduces or increases the pulsation rate (Figures 3D, 3F, 4E, and 4F). Importantly, changes in the frequency of run and tumble transitions are also regulated by the signaling level of the transmembrane receptor Cxcr4b, which directs the cells and provides them with information on their distance from the target.^19^ The control over pulsation features is thereby important for migration precision (Figure S8C, upper panels); if an increase in pulsation frequency does not occur when the cells approach the developing gonad, the run phases can be too long, causing cells to pass by their target and arrive at ectopic locations (Figure S8C, middle panels).^19^ Conversely, if the transition frequency between run and tumble phases is too high, the cells will spend more time in the immotile phase, thereby having a slower overall migration speed, resulting in cell localization at ectopic sites (Figure S8C, bottom panels).^25,26,85,86^

Combining our data with recent results,^20^ we present a model that accounts for the generation of, and control over, run and tumble behavior (Figure S8A and 8B). During run phases, membrane-bound active Prex1 interacts with and activates Rac1, which, in turn, promotes actin polymerization and, with a certain time delay, activation of Pak1. The kinase then phosphorylates and thereby inhibits Prex1 function. The inhibition of Prex1 reduces Rac1 activity, leading to a depletion of polymerized actin and the concomitant loss of the cell front (Figure S8B, Step 1). Therefore, we assume that the entry into the tumble phases results from low activity of Prex1 and Rac1 (Figure 1C, 1D, and 1G) mediated by the activation of Pak. Under such conditions, the loss of actomyosin-based contractility and retrograde flow is also manifested by the homogeneous distribution of linker proteins (such as Ezrin and Esyt2a) at the cell rear (Figure S8B, Step 2, see also Figure 1E, 1F, and 1G).^20^ The activation of Prex1 and the ensuing Rac1 activation and actin polymerization can, therefore, be considered the initial step by which cells exit the tumble phase (Figure S8B, Step 3). The activation of Rac1 and, thus, the position where the front is defined can be either stochastic around the cell in the absence of a polarized directional signal, or biased by polarized activation of Cxcr4b by its ligand Cxcl12a, thereby directing the next run toward the attractant source.^20^ Following the polarized accumulation of actin, retrograde flow defines the non-blebbing rear, ensuring robust cell polarity and effective forward migration during run phases (Figure S8B, Step 4).^20^

In zebrafish PGCs, the transitions between run and tumble phases themselves are cell-autonomous. However, in addition to steering the cells, Cxcl12a can also affect the duration of run and tumble phases.^18^ By activating G*β*γ subunits, an upstream regulator of Prex1 activity,^35,37^ Cxcr4b signaling could modulate the feedback loop. Consistently, upon knockdown of G*β*γ signaling, zebrafish PGCs fail to polarize, and exhibit reduced Rac1 activity.^86^ In addition to G*β*γ, Prex1 is activated by PtdIns(3,4,5)P^3^ (PIP^3^).^35,87^ However, in contrast to other cell types, in zebrafish PGCs PIP^3^ is uniformly distributed around the cell membrane.^85^ Thus, PIP^3^ could act as an apolar regulator of Rac1 activity, while G*β*γ signaling could function downstream of Cxcr4b to bias the initial activation of Prex.

We also found that the position of the cell front in subsequent run phases was biased in cdPGCs, despite being separated by seemingly apolar cdTumble phases (Figure 7). Such behavior provides evidence of a molecular memory, which has been reported in other cell types such as migrating leukocytes,^13^ *Dictyostelium*^88^ and neutrophils.^63,89^ Importantly, the significant bias we detected in the position of successive cell fronts in cdPGCs is diminished in non-manipulated PGCs that form blebs. This raises the possibility that apolar blebbing during tumble phases may serve to reduce inhomogeneities in the distribution of polarity-inducing components (Figure 7D). Indeed, the long-lived asymmetric distribution of such components is thought to be the basis for directional memory in the migration of other cell types.^63,64^ According to this hypothesis, factors that facilitate Rac1 activation and, thus, play a role in maintaining cell polarity during run phases are dispersed by blebs during tumble phases. Such a mechanism could allow a “search” migration mode in the absence of a guidance cue, as well as an efficient response to changes in directional information.

The molecular networks that regulate oscillations and their biological functions are largely unknown. Our results show that pulsations in polarized Rac1 activity can be regulated by a module of just a few proteins and, importantly, we also identify molecules and processes that are dispensable for this process. This means that the cells possess an intrinsic pacemaker-like molecular network that initiates polar pulsations. The pulsations, in turn, control the transitions between run and tumble states during germ cell migration. Considering that many cell types display a similar behavior as they introduce corrections to their migration path, molecular networks similar to the one identified here are likely important and widespread in other cell migration processes.

## Acknowledgements

We thank Celeste Brennecka for editing and Michal Reichman-Fried for critical comments on the manuscript. We thank Ursula Jordan, Esther Messerschmidt and Ines Sandbote for technical assistance. We thank Jan Schick for designing the cartoon models of zebrafish embryos. We thank Aneta Koseska for a constructive discussion. N.S.G. is the incumbent of the Lee and William Abramowitz Professorial Chair of Biophysics (Weizmann Institute), and acknowledges support from the Royal Society Wolfson Visiting Fellowship.

D.H., L.K. and E.R. are supported by funding from the University of Muenster, the Max Planck Institute for Molecular Biomedicine and the German Research Foundation (DFG) grant RA 863/14-1 and SFB 1348 project B06. N.S.G. is the incumbent of the Lee and William Abramowitz Professorial Chair of Biophysics (Weizmann Institute), and acknowledges support by the Royal Society Wolfson Visiting Fellowship. B.D.S. and T.A. are supported by the Wellcome Trust (219478/Z/19/Z) and B.D.S. by a Royal Society EP Abraham Research Professorship (RP/R1/180165 and RP\R\231004).

## Author contributions

E.R. supervised the project; D.H. and E.R. designed the work; D.H., E.R. and T.A. wrote the manuscript; and D.H. performed all the experiments; T.A., N.G. and B.S. designed the mathematical model; L.K. analyzed the angle of consecutive polarity vectors and prepared the rose plots. All authors read and approved the final manuscript with minor revisions.

## STAR Methods

## Resource availability Lead contact

Any information or requests for resources and reagents should be directed to the lead contact, Erez Raz.

## Materials availability

All plasmids generated in this study are available from the lead contact.

## Data and code availability

- No large-scale datasets have been generated in this study. Microscopy data reported in this paper will be shared by the lead contact upon request. Accession numbers are listed in the key resources table.
- This paper does not report original code.
- Any additional information regarding the mathematical model is available from the corresponding author, Tal Agranov.
- Any additional information required to reanalyze the data reported in this paper is available from the lead contact upon request.

### Experimental Model and Study Participant Details Zebrafish strains and maintenance

The following zebrafish (*Danio rerio*) lines were used in this work: wild-type strain AB genetic background, wild-type strain of AB/TL genetic background and the transgenic lines Tg(*kop:lifeact-egfp-nos3’UTR-cry:DsRed*),^90^ Tg(*kop*:*mCherry-f-nos3’UTR-cmlc:egfp*),^91^ Tg(*kop:egfp-f-nos3’UTR-cry:DsRed*).^26^ The fish were maintained according to the recommendations listed in 2007/526/EC and Article 33 of European Union Directive 2010/63. Maintenance was performed in compliance with the German, North-Rhine-Westphalia state law, following the regulations of the *Landesamt für Natur, Umwelt und Verbraucherschutz Nordrhein-Westfalen* and was supervised by the veterinarian office of the city of Muenster.

## Method Details

### Plasmid cloning, mRNA synthesis and embryo microinjection

Two nanoliters of sense mRNA and/or translation-blocking morpholino antisense oligonucleotides (MO, Genetools) were microinjected into the yolk of 1-cell stage embryos. mRNA was synthesized using the mMessage mMachine SP6 or T3 transcription kit (Thermo Fisher Scientific). For cloning of plasmid constructs, a restriction-free cloning approach^92^ was employed. Mutations within the sequence were induced via PCR-based site-directed mutagenesis approach adapted from the QuickChange™ site-directed mutagenesis protocol with either overlapping^93^ or non-overlapping primers.^94^ For this work, the full-length zebrafish Prex1 was amplified from cDNA and cloned into a pSP64 vector. Dominant-negative (DN) Prex1 was generated by introducing the mutations E35A and N215A (based on Hill et al. (2005),^37^ homologous to the mutation in human Prex1 E56A/N238A). For Prex1 ΔDH, amino acids (aa) 28 - 217 were deleted (based on Yoshizawa et al. (2005),^38^ the homologous deletion in human Prex1 comprises aa 49 - 240). Prex1 PH/DH domains (aa 1 - 368) were amplified from cDNA and used as the constitutively active (CA) form (based on Hill et al. (2005),^37^ the homologous domains in human Prex1 comprise aa 1 - 396). The DN zebrafish Pak1 K331R mutation was generated based on the human mutation K299R^45^ and the CA Pak1 T455E mutation was based on the human mutation T423E.^46^ For the generation of the constitutively active and Rac1-binding-deficient Pak1, the mutations H82L and H85L were induced (based on Sells et al. (1997),^46^ H83L/H86L in humans). For protein expression specifically in PGCs, the corresponding open reading frames (ORFs) were cloned upstream of the *nanos3 3’UTR* sequence,^95^ while for expression in all cells (somatic and PGCs) the 3’UTR sequence of the *Xenopus globin* gene was included downstream of the ORFs. A list of mRNAs, morpholinos, and the respective concentrations used in this study are provided in the supplementary information, Table S1. Embryos were kept in 0.3× Danieau’s solution and incubated at 25°C or 28 °C, depending on the experiment.

### Imaging

Embryos were dechorionated prior to image acquisition, transferred to 1.5 % agarose ramps covered with Danieau’s solution and maintained at 28 °C during imaging. Embryos at 24 hpf were anesthetized using 0.64 mM Tricaine (Sigma-Aldrich) in 0.3x Danieau’s solution prior to imaging. Spinning disk microscopy was conducted using Carl Zeiss Axio Imager Z1 and M1 microscopes equipped with Yokogawa CSUX1FW-06P-01 spinning disk units. Images and time-lapse movies were acquired using a Hamamatsu ORCA-Flash4.0 LT C11440 camera and Visitron Systems acquisition software (VisiView). Imaging of actin dynamics was conducted between 7 - 10 hpf using a 40x or 63x objective. In all experiments, time-lapse movies were captured with an 8 sec time interval between frames and 20 µm Z-stacks (5 Z-planes, each 5 µm apart). 24 hpf embryos were imaged with a 5x objective. 200 µm Z-stacks were acquired (21 Z-planes, each 10 µm apart). Confocal laser scanning microscopy was performed using a LSM710 (Zeiss) upright microscope and ZEN software (Zeiss) and a 40x objective.

### Analysis of ectopic PGCs

To quantify the number of PGCs located in ectopic positions, Tg(*kop:lifeact-egfp*-*nos3’UTR*- *cry:DsRed*) embryos were used for Figure 5A and 5B and Tg(*kop:egfp-f-nos3’UTR-cry:dsRed*) embryos were used for Figures 5D and S3A. PGCs were counted on one side of the embryo using a 10x objective. Ectopic PGCs were defined as germ cells located outside the gonad region, which starts at the anterior border of the yolk extension toward the posterior, until the middle of the yolk extension. The ectopicity ratio was calculated by dividing the number of ectopic cells by the total number of germ cells in each embryo.

### Derivation and analysis of the mathematical model

Here, we detail further the derivation of the minimal mathematical model that we used to study the phase behavior and dynamics of the polar pulsations.

#### Biochemical reaction modeling

To develop a minimal modelling scheme, we placed emphasis on just three chemical components, Prex1, Rac1 and Pak1. For simplicity, we assumed that the inactive protein species are kept at roughly constant bulk cytoplasmic concentrations. Indeed, this assumption is supported by prior studies that explored the behavior of these species in related biological contexts.^55,96,97^ However, our qualitative findings are in any case largely insensitive to this assumption provided spatiotemporal variations in the cytoplasmic concentration are small compared to those at the membrane surface of the cell. This holds true when the local concentration of the inactive bulk population exceeds that of the active population and is therefore affected little when a fraction of the molecules is recruited into the membrane. At the same time, the cytoplasmic components are expected to remain roughly homogeneous due to fast bulk diffusion. If the population of activated molecules that are recruited to the membrane surface is sufficiently large that the local concentration becomes comparable to the bulk cytoplasmic population, then the model would need to be revised so that total number conservation of molecules is taken into account. We note that such effects play a crucial role in other systems displaying wave pinning.^98^

Based on the proposed regulatory network (Figure 6A), the dynamics of active Prex1, Pak1 and Rac1 concentrations are defined by the coupled reaction-diffusion system,

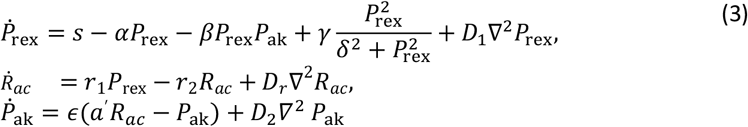

Here, the terms linear in *r*_1_and *a*′ represent, respectively, the activation rates of Rac and Pak1 on the membrane. Here, we assume that the local activation rate, per unit surface area across the membrane, is proportional to the product of the bulk concentration of the inactive species ρ_cyto_ and the surface number density of the active species *n*_active_. A 1:1 stoichiometry was described for the interaction between Prex1 and Rac1.^99^ Similarly, Rac1/Cdc42 is presumed to activate Pak1 dimers with a 1:1 stoichiometry,^56^ though evidence also supports an alternative 1:2 stoichiometry (Rac1-Pak1) in some contexts.^100^ The stoichiometry between Pak1 and Prex1 during the phosphorylation reaction and inactivation of the GEF is unknown, but will be assumed in the model to be 1:1 (see below). Then since the cytoplasmic bulk concentrations are assumed roughly constant, these rates are linear in the surface number density of active species. Notice that Prex1 is also activated by upstream membrane-bound regulators,^35^ which in the absence of external cues are assumed to be uniformly distributed at some constant number density across the membrane. This then leads to a constant and uniform activation rate for Prex1 given by *s*. The local deactivation rate of Prex1 is represented by the term −*βP*_ref_*P*_ak_, with this reaction involving one active Pak1 and one active Prex1 molecule^1^. The natural deactivation terms −*αP*_ref_, −*r*_2_*R*_ac_ and −ϵ*P*_ak_ ensure the condition that all three species are stable in their inactive form. As Pak1 is reported to have a slow response, as suggested by the slow dissociation constant of active Rac1 and Pak1,^56^ and the temporal profiles of Pak1 activation^101^ and of Prex1 inhibition by Pak1,^43^ ϵ is smaller compared to other rates, and this will be important in the following. Note that, as the Prex1 dynamics have an additional nonlinear deactivation rate −*βP*_ref_*P*_ak_, one can omit the linear term −*αP*_ref_ without affecting the qualitative behavior of the system^2^. Therefore, since these equations are more tractable without this term, in the following it is omitted. Without further nonlinearities, the dynamics of the model would admit a single *stable* fixed point at all range of parameters. The pulsating behavior observed in experiment indicates the existence of an additional amplification, which we include as an autocatalytic term in the Prex1 dynamics. We model it here as a Hill-type function with coefficient 2. However, once again, our qualitative findings are insensitive to the precise form of the amplification within some mild requirements discussed below.

The reaction dynamics are supplemented by diffusion terms. Here, we consider a simple one-dimensional ring geometry to model the cell membrane where, without loss of generality, we set the circumference to unity (absorbing the space dimensions into a rescaling of the effective diffusion coefficients). Further, to map the phase behavior of the model analytically, we will also consider an even simpler two-site geometry mimicking a single axis of potential polarization (Figure 6B).

To proceed, it is helpful to further rescale the three concentration fields and time,

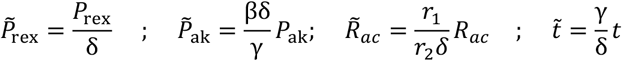

which identifies just 4 effective parameters from the original eight. In this case, the reaction-diffusion equations take the form

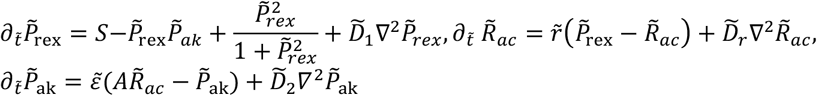

with the dimensionless reaction parameters and rescaled diffusion coefficients 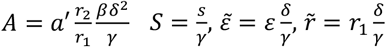, and 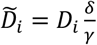 with *i* = 1,2, *r*. In the following, we omit the tilde for ease of notation.

#### FitzHugh–Nagumo nature of the dynamics

The simple linear nature of the Rac1 dynamics means that the qualitative behavior of the model dynamics (3), and in particular the set and stability of homogenous fixed points^3^, are entirely dominated by Prex1 and Pak1 concentration. In this case, the dynamics can be captured by considering a reduced set of equations obtained by “adiabatically” eliminating Rac1, with R_*ac*_ ≈ *P*_ref_, from which we obtain the reduced set of equations,

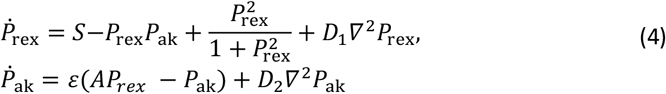

These equations represent the rescaled version of the model presented in the main text (1) with *a*′ = *ar*_1_/*r*_2_. Formally, this reduction only holds in the limit of fast Rac1 dynamics *r* ≫ 1. However, numerically, we find that the qualitative behavior also holds for moderate values *r*, particularly in the regime of slow Pak1 dynamics ϵ ≪ 1, which we are interested in here.

The set of equations (4) has the general structure of a FitzHugh–Nagumo type system with Prex1 and Pak1 taking the role of the excitatory and inhibitory species, respectively.^48,49,102^ This is evident when looking at the reaction term nullclines for Prex1 and Pak1, respectively (Figure 6D–G)

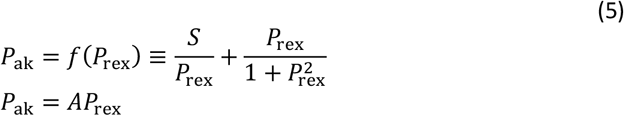

The two nullclines always intersect at a single fixed point,

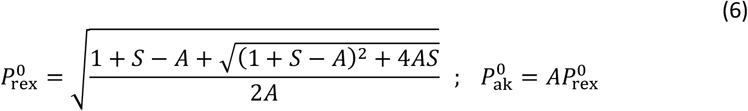

In the range 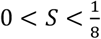,the Prex1 nullcline *f* (*P*_*ref*_) has the signature of an (inverted) N-shaped curve with a single minima and maxima. This feature sets the phenomenology that we now discuss, and it arises due to the self-enhancement Hill-type term that we included for the Prex1 dynamics. Importantly, this feature is not particular to the Hill-type term and will be generic for any nonlinear self-enhancement that saturates at large abundance.

The set of equations (4) have featured in prior works studying polarity in different types of chemotactic cells.^58,81^ However, these studies considered a different regime of parameters that leads to different dynamical behaviors then those we report on here. In particular, as shown in the following, the regime of polar pulsations observed here emerges when the differential in the diffusion coefficients is sufficiently large,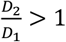. If this condition is not met, the system of equations (4) exhibit homogeneous oscillations and propagating waves, which were the focus of previous works.^57,58^

To gain insight into the dynamics (4), we start with an analysis of the zero-dimensional model (where the active protein concentrations are space-independent), then consider a simplified, yet analytically tractable, two-compartment model, before moving to the more realistic one-dimensional ring geometry.

#### Dynamics of the zero-dimensional case

In the absence of diffusion, the zero-dimensional model represents a simple version of a FitzHugh–Nagumo type system. Its behavior can be described by considering a fixed value of *S* in the range 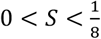 and varying *A*. At very large or small *A* the fixed point (5) lies to the left of the Prex1 nullcline minima, or the right of its maxima, respectively (Figure 6D and 6G), where it shows a stable fixed point. Within an intermediate range of *A* values, it loses stability, turning into an unstable spiral. This happens when

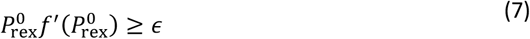

where the prime denotes the derivative with respect to the argument. At small enough ϵ, the equality is always satisfied at two instances when the fixed point is located within the middle branch of the Prex1 nullcline, between the two extrema. At vanishing ϵ, these locations approach the two extrema. These two points mark a pair of super-critical Hopf bifurcations that engulf a phase of a limit cycle that grows continuously from the stable fixed point. When *A* is large such that the fixed point is stable, but approaching the Hopf bifurcation point (7), the system displays the signature excitatory behavior (Figure 6E).^59,61^

#### Complete analytical mapping of the dynamic phase behavior for a simplified two-compartment model

To understand the phase behavior of the spatially extended system, we start by considering a simplified two-compartment model, where we denote by + and – the two spatial compartments coupled by Pak1 diffusion

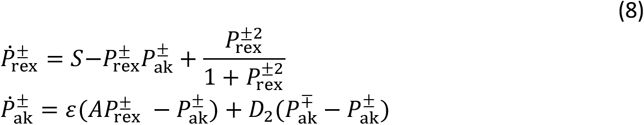

Here, since bound Prex1 is expected to have a much slower diffusion, we set the effective Prex1 diffusion coefficient to zero, when the equations become analytical tractable. (Yet, we have verified numerically that the qualitative behavior holds when the Prex1 diffusion coefficient is non-zero.) The complete phase behavior is presented in Figure 6B and 6C. The analysis starts with a straightforward linear stability analysis which reveals that, in addition to the Hopf bifurcation (7), the homogenous fixed point 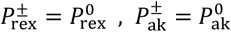 can now also lose stability due to a Turing-type instability. This happens when

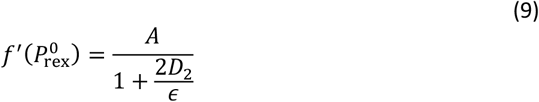

As for the Hopf bifurcation, this condition is satisfied in two instances when the fixed point is located within the middle branch of the Prex1 nullclines (a situation guaranteed at small ϵ and any non-zero value of the diffusion coefficient *D*_2_). These coincide with the two nullcline’s extrema at vanishing *ϵ*. These two points are marked by green dots in Figure 6B and 6C. Importantly, at sufficiently large *D*_2_, these two Turing-like instabilities *engulf* the two Hopf bifurcation points given by (7). At small *ϵ* this happens as soon as

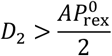

In this case, a pitchfork bifurcation precedes the Hopf bifurcation giving way to stationary “polar” states with 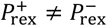.One of the pitchfork bifurcations is super-critical and the other is sub-critical (Figure 6C). The steady-state values of 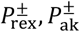 at this emerging symmetry breaking polar state can be depicted graphically by the intersection of two shifted Pak1 nullclines (marked by red dashed lines in Figure 6F–H) with the Prex1 nullcline. Indeed, since we set the Prex1 diffusion coefficient to vanish, then the local composition of the two compartments must still lie on the Prex1 nullcline (5). These shifted nullclines are given by

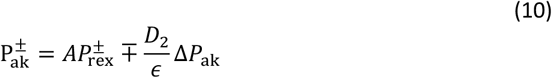

with 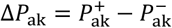 determined self-consistently by solving for the intersection with the Prex1 nullcline. Following some steps of algebra, one finds two branches of solutions

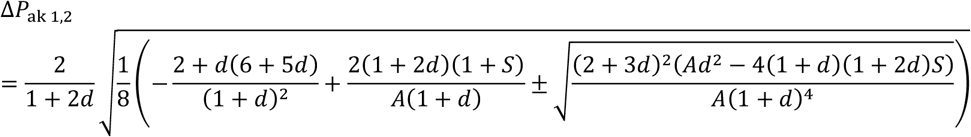

with

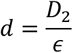

which correspond to the two pitchfork bifurcations. One of them is a branch of unstable polar states, while the other is (in parts of the phase diagram) stable (black lines in Figure 6C). Following some more algebra, this provides the local composition

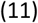

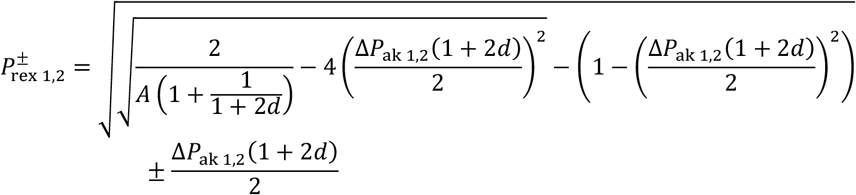

and 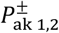 follows through equation (10). Importantly, one can determine the stability of this polar state and find that it is subject to a pair of super-critical Hopf bifurcations marking the onset of polarity pulsations (see red lines and dots in Figure 6^4^). These Hopf bifurcations are different from (7), which correspond to the homogeneous state, but are closely related to them. Specifically, if the local composition in one of the compartments lands on the middle branch of the Prex1 nullcline (which is subject to the Hopf instability (7)), then the composite polar state loses stability and undergoes a Hopf bifurcation (Figure 6F). This simplified local analysis becomes exact at vanishing *ϵ*, where the unstable mode that corresponds to the Hopf bifurcation has a component only along the Prex1 axis and is not affected by the Pak1 diffusive coupling. Conversely, if the local composition of both compartments lands outside the middle branch, they are both locally stable and now comprise a stable polar state, which may or may not coexist with a stable apolar state (Figure 6G and 6H). This behavior is summarized by the phase diagram and bifurcation diagram along a fixed *S* cross-section (Figure 6B and 6C). Using the simple two-compartment model as a guide, we now turn to the phase behavior of the continuous one-dimensional model, which we find to be qualitatively similar.

#### Continuous one-dimensional model

Here, we are only able to analytically locate the Turing and Hopf instabilities about a homogenous state. The Hopf bifurcation condition (7) remains the same as it only involves homogenous perturbations. The Turing instability (9) is modified. Applying a linear stability analysis of equations (4) around the steady state protein concentrations 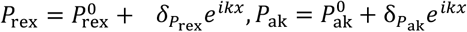, we find that the corresponding stability matrix

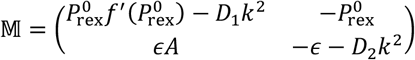

Since all components of 𝕄 are real, a Turing instability sets in when both the trace of 𝕄 and its determinant becomes negative, i.e.

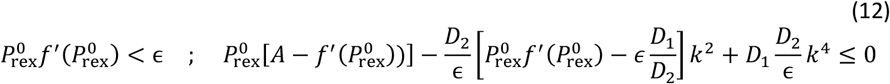

respectively. The determinant is parabolic in *k*^2^with the constant and the quadratic coefficients strictly positive 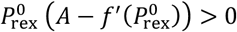 and 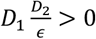 for all parameter values.

Thus, the determinant becomes negative when the linear coefficient is sufficiently negative. Combined with the condition on the trace, this happens when

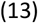

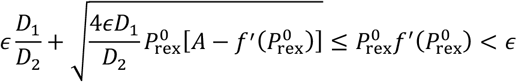

This condition can always be realized at a sufficiently large diffusion coefficient disparity. At small *ϵ* this requires

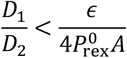

The first instability therefore occurs at a wave number given by the condition

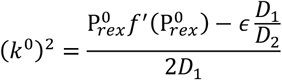

Depending on the system size, this unstable mode might or might not fit within the system. In the latter case, linear instability initiates when the determinant (12), evaluated at the principal mode *k* = 2π, becomes negative. This will modify the instability condition (13). In either case, the instability can be sub-critical, and we did not attempt to classify its type. In our numerical simulations of the dynamics (4), an instability appeared when the left equality in (13) was satisfied. In addition, the final patterns that emerged spanned the entire system size. In particular, we did not observe Turing patterns comprising multiple peaks. The latter might arise in longer systems (i.e. for smaller values of the re-scaled diffusion coefficients), though we do not explore this possibility here. The corresponding phase behavior, summarized in Figure 6, is qualitatively the same as in the two-compartment geometry (see corresponding parameter values in the caption). At large negative feedback *A*, the system lies in the excitatory regime (Figure 6E, bottom panel). Lowering the value of *A* below the Turing instability, we observed two possible transitions. For not too small values of *ϵ*, we observe the same phase behavior as for the two compartment model: first a transition into a polar state, before a Hopf-type instability mediates a transition into polar pulsations (Figure 6F, bottom panel). However, in contrast to the two-compartment model, for very low values of *ϵ*, we observed a direct, discontinuous, transition into polar pulsations, skipping the intermediate polar regime. Nevertheless, it might be that the range of parameters stabilizing this intermediate polar regime is very narrow and hard to discern in simulations. Since this intermediate region is in any case small, it does not impact on our conclusions. Upon further lowering of *A*, a stable polar pattern emerges (Figure 6G, bottom panel). Lastly, at values of *A* that now pass the second Turing instability point given by equation (13), the system displays stable apolar states (Figure 6H, bottom panel).

#### Comparison with experiment

The polar pulsation regime (Figure 6F) agrees well with the experimental observations of cdPGCs. In particular, we found that the pulsation behavior of the model is robust with respect to noise (Figure 6F, bottom panel), where polar runs display transient directional memory. The model also nicely accounts for the experimental Pak1-treated cells shown in Figure 4. The DN Pak1 treated cells overexpress a form of Pak1 that does not deactivate Prex1. As only a portion of the Pak1 molecules now deactivate Prex1, this then decreases the deactivation rate *β*, which would decrease the value of the control parameter *A*. In this case the phase diagram (Figure 6B) would predict a shift towards the stable polar state in good agreement with experimental observation. This is evident in Figures 4B and S7, where, isolating the cells that were strongly affected by the treatment (cells with run phases >10 min), one observes a persistent static polar amplitude that did not decrease throughout the duration of the imaging (Figure S7B). A similar effect occurs under the Rac1^C40^ treatment. Indeed, this form of Rac1 cannot activate Pak1, leading to a reduced activation rate *a* and a subsequent reduction of the control parameter *A*. Again, the experimental observation agrees well with a stable polar regime (Figure 4H). Lastly, the CA Pak1 treatment means that there is a population of stably active Pak1 molecules. Let us denote their effective surface number density by *C*. These should affect the Prex1 deactivation term −*βP*_ak_*P*_ref_ → −*β*(*P*_ak_ + *C*)*P*_ref_, which in turn would lead to a downward shift of the Prex1 nullcline (5) *P*_ak_ = *f*(*P*_ref_) → *P*_ak_ = *f*(*P*_ref_) − *C*. In this case, the nullclines intersect in the excitatory branch. Indeed, the experimental observations agree with the excitatory dynamics of rapid runs separated by broadly distributed tumbles. This is evident in Figures 4F, S4A and S6, where, isolating the cells that were strongly affected by the treatment, one observes an irregular pattern of many frequent but modest amplitude runs, with an occasional more protracted polar run (Figure S6).

The Prex1 treated cells (Figure 3) also fall within the model’s dynamical regimes. The hyperactive Prex1 cells have an elevated effective activation rate *S*. This pushes the system upwards in the phase diagram (Figure 6B) into the stable polar regime, in nice agreement with the experimental observation (Figure 3D). The constitutively active Prex1 treated cells have a population of stably active Prex1 molecules, with a surface area we denote by *C*. This pushes the Pak1 nullcline upwards *P*_ak_ = *aP*_ref_ → *P*_ak_ = *aP*_ref_ + *C* causing the nullclines to intersect in the excitatory branch. Yet the actin polymerization now is dominated by the population of stably active Prex1 molecules, rather than the occasional modest excitations of the stably inactive population. The former is homogenously distributed across the cell, and this agrees with experimental observation (Figure 3G).

### Quantification and Statistical Analysis

#### Quantification of cell polarity

The polarity value was calculated using an intensity-based Matlab (Mathworks) program that follows the approach described in Olguin-Olguin.^20^ Initially, a mask covering the complete shape of the cell was generated by thresholding using Fiji, and the cell center (COM) was determined by averaging over the positions of all pixels inside the cell for each frame. The center of each fluorescence (COF) signal was then determined by a weighted average over all pixel positions using the fluorescence intensity as weight and normalizing by the sum of all fluorescence intensities. The vector from the COM to COF gives a polarity vector. A scalar polarization value can be determined by projecting the fluorescence intensity of each pixel on the polarization vector. Briefly, the cosine of the angle between the polarity vector and the individual vector of each pixel was multiplied by the intensity of the respective pixel. The sum of these projected values was then divided by the total sum of the fluorescence signal in the cell. The following equation was used:

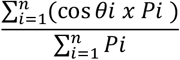

The polarity values range from 0.0 for no polarization (homogeneous fluorescence intensity) to 1.0 for full polarization (all fluorescence concentrated in one pixel).

A threshold of 0.1 was set to distinguish polarized run phases (> 0.1) from apolar tumble phases (< 0.1). In addition, to be considered a run or a tumble phase, three consecutive time points (i.e. 24 sec) were required to cross the threshold value. To be defined as a run phase, at least one time point needed to exceed a polarity value of 0.15.

To analyze the polarity value of Rac1 GTP (Pak1 GBD) in contractility-deficient PGCs (cdPGCs), the contrast was increased using the Fiji software.

To average the polarity values during run to tumble transition for multiple PGCs, the depolarization start point of actin polarity was defined as t = 0 min. The depolarization start point was defined as the peak value preceding a continuous decrease in the polarity value, until the value was lower than the 0.1 threshold.

To average the polarity values from multiple PGCs over 10 min (e.g. Figure 2D) the middle of the run phase was considered as t = 0 min. The polarity values of the preceding tumble phases were plotted up to 5 minutes before and after the t= 0 min point. For cells that exhibit run or tumble phases longer than 10 minutes, an arbitrary point was designated as the middle.

### Quantification of angles between polarity vectors of consecutive run phases

The direction of the cell front during run phases was determined by the polarity vector (see Quantification of cell polarity section). To quantify the angle between the cell fronts of consecutive run phases, the direction of the polarity vector was measured with Fiji at two time points: (1) during the last time frame (8 sec intervals) of the first run phase, before depolarization and transition into a tumble phase, and (2) during the first time point of the new run phase. Then, the angle between these vectors was quantified.

### Rac1 FRET analysis

Imaging and analysis for the Rac1-FRET were performed at 8 - 10 hpf on an LSM710 confocal microscope (40x, numerical aperture (NA) 0.75, pinhole 10 mm, 512 × 512) controlled by ZEN software (Zeiss). Single images of one focal plane were prepared. Analysis was done using ImageJ software (the protocol was adapted from Tarbashevich et al.^91^). The Rac1 FRET values were normalized to the average measured values of the control cells. To quantify the Rac1 FRET values specifically during run and tumble phases, the Rac1 FRET values measured during run and tumble phases, respectively, were normalized only to the average values of the respective phase of the control cells.

### Quantification of bleb frequency

PGCs were imaged between 7 - 10 hpf using a spinning disc confocal microscope (63x objective, 10 min at 8 sec intervals). The bleb frequency was quantified by counting the number of blebs and dividing this number by the analyzed period of time.

### Analysis of run and tumble phase durations

PGCs were imaged between 7 - 10 hpf using a spinning disc confocal microscope (63x objective, 10 - 30 min at 8 sec intervals). Phase durations were considered only if the beginning and end of the respective phases were within the imaged time period, except for run phases longer than 10 min. Run and tumble phases were defined based on the quantified polarity values (see above).

### Polarized cell ratio analysis

PGCs were imaged between 7 - 10 hpf using a spinning disc confocal microscope (63x objective, 10 min at 8 sec intervals). Initially, tumbling cells were imaged for 10 min. PGCs that transitioned to a run phase were considered polarized cells. To calculate the ratio of polarized cells, the number of polarized cells was divided by the total number of imaged cells. Run and tumble phases were defined based on the polarity values quantified as described above.

### Image processing

Time-lapse images were analyzed and edited using the Fiji software with adjustments of brightness and contrast. Background subtraction (radius = 200 pixels (px) for Lifeact, Ezrin and Esyt2a and radius = 25 px for Pak1 GBD) and bleach correction (Simple Ratio) was conducted using Fiji. Maximum intensity projections (MIPs) were generated from Z-stack images in Figures 2B, 3A, 3D, 3G, 4B, 4E, 4h, 5A, 5D, 7A, 7B, S2C–F, S3A, and S5A.

### Statistics

Statistical analysis and graph creation was conducted using the GraphPad Prism software. The Rose plots (Figure 7C) were designed with the python-based package ‘matplotlib’ (Version 3.9.2). The statistical tests used, the n and the P-values are indicated in the graphs, figures and legends. All experiments were repeated at least three times.

## Supplemental Information

Document S1. Figures S1–S8 Table S1. List of injected mRNAs and morpholinos, Related to STAR Methods Video S1. Dynamics of actin distribution during run and tumble phases, related to Figure 1 Video S2. Dynamics of actin and Rac1 GTP distribution during run to tumble transition, related to Figure 1 Video S3. Dynamics of actin and Esyt2a distribution during run to tumble transition, related to Figure 1 Video S4. Polar pulsations of actin in a cdPGC, related to Figure 2 Video S5. Dynamics of Ezrin during polar pulsations of actin in a cdPGC, related to Figure 2 Video S6. Polar pulsations of actin and Rac1 GTP in a cdPGC, related to Figure 2 Video S7. Dynamics of actin distribution in a PGC and a cdPGC expressing DN Rac1, related to Figure S2 Video S8. Dynamics of actin distribution in a PGC and a cdPGC expressing CA Rac1, related to Figure S2 Video S9. Dynamics of actin distribution in a cdPGC expressing DN Prex1, related to Figure 3 Video S10. Dynamics of actin distribution in a cdPGC overexpressing Prex1, related to Figure 3 Video S11. Dynamics of actin distribution in a cdPGC expressing CA Prex1, related to Figure 3 Video S12. Dynamics of actin distribution in a cdPGC expressing DN Pak1, related to Figure 4 Video S13. Dynamics of actin distribution in a cdPGC expressing CA Pak1, related to Figure 4 Video S14. Dynamics of actin distribution in a cdPGC expressing Rac1^C40^, related to Figure 4 Video S15. Dynamics of actin distribution in a cdPGC overexpressing Rac1, related to Figure S5

## Footnotes

1 The actual stoichiometry is unknown. Adopting an alternative stoichiometry of 2:1 (Pak1-Prex1) will not affect the qualitative behavior of the system much since the shape and relative positioning of the reaction rate nullclines remains the same, see below.

2 However, at large enough deactivation rate *α*, the number of homogeneous fixed points can increase, with possible bi-stability. We do not attempt to explore the corresponding phase behavior here.

3 In fact, at vanishing *ϵ*, the unstable eigenvalue that corresponds to homogeneous perturbations about the fixed point of (1), coincides with that of the reduced set (4). This calculation is not presented here.

4 For this, one has to compute the spectrum of the four-by-four stability matrix and show that a pair of complex eigenvalues crosses the imaginary axis. This can be done analytically using the Routh–Hurwitz criterion. We found it simpler to evaluate the spectrum numerically.

