## Supplemental information for "Corrections in single cell migration path in vivo are controlled by pulses in polar Rac1 activation"

**Hoffmann et al**

### Supplemental Information

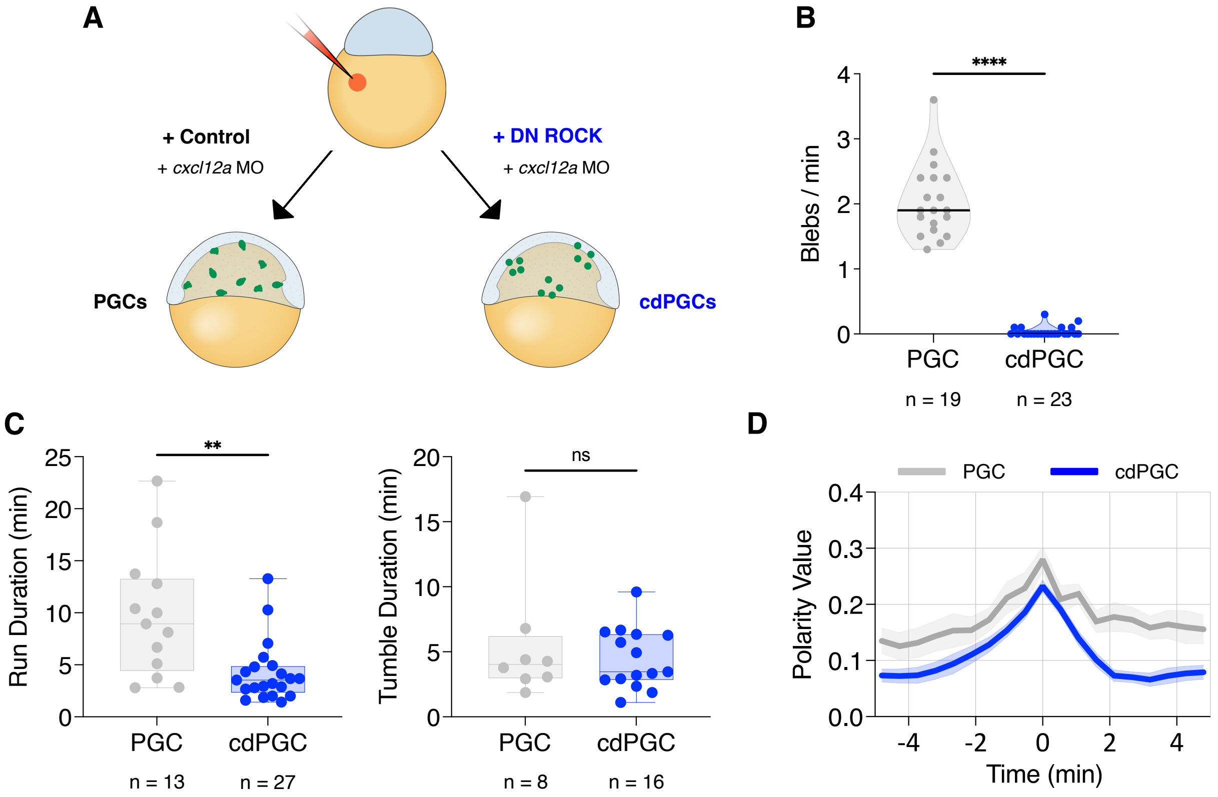


**Figure S1. Comparison between control and contractility-deficient PGCs (cdPGCs), Related to Figure 2**

(A) Illustration showing the procedure for knocking down contractility for generating cdPGCs by injection of a dominant-negative (DN) version of ROCK. Cells expressing a control protein were used as control PGCs. The red color represents the injection mix that contains anti-*cxcl12a* morpholinos (MO), and the green dots represent PGCs (left) or cdPGCs (right).
(B) The number of blebs formed per minute in control PGCs and cdPGCs. ****, P-value < 0.0001 (Mann-Whitney test).
(C) Duration of run (left) and tumble phases (right) in control PGCs and cdPGCs. **, P-value = 0.0019; ns = non-significant (Mann-Whitney test).
(D) Average polarity value (see STAR Methods) observed in control PGCs (grey) and cdPGCs (blue) over time with respect to the median time point of the respective run phase (time 0). Shaded areas represent the SEM and n is the number of cells analyzed.

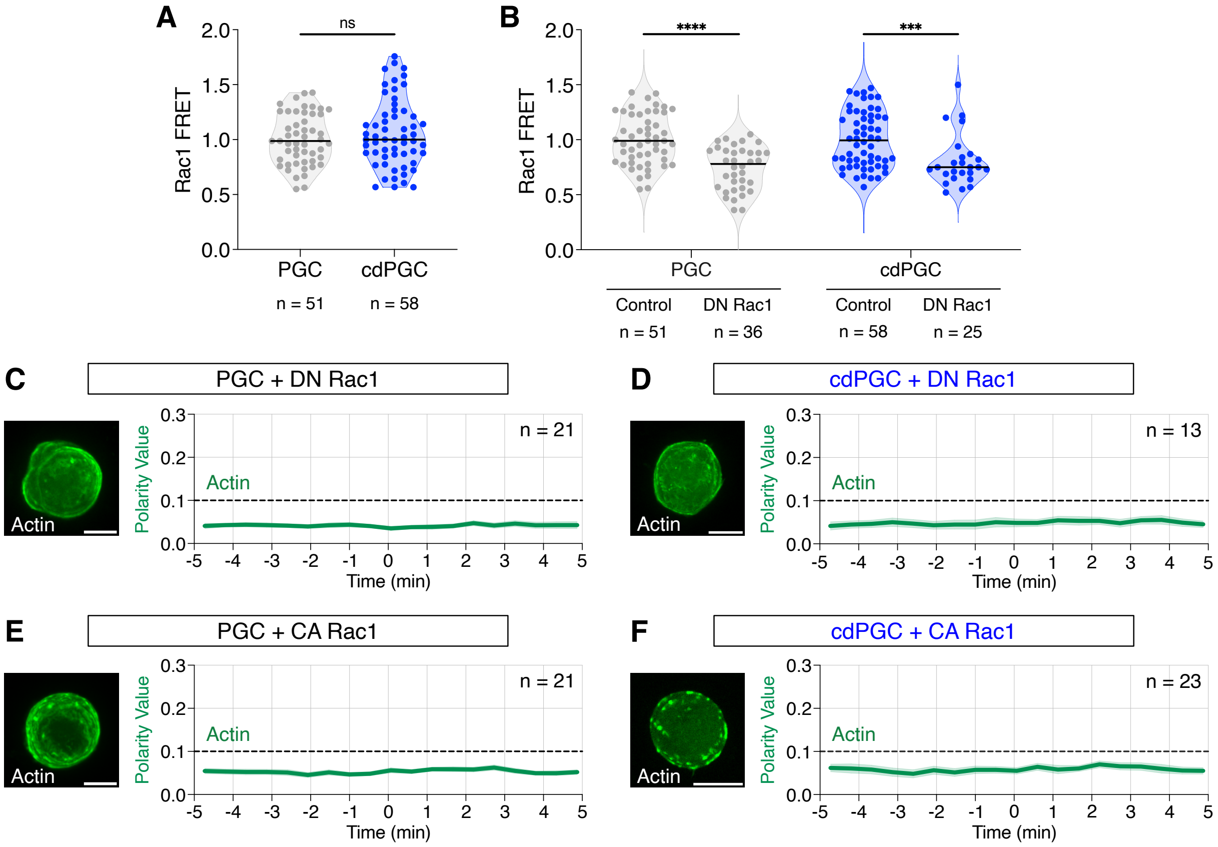


**Figure S2. Manipulation of Rac1 activity in PGCs and cdPGCs, Related to Figure 2**

(A) Rac1 FRET ratios for control PGCs and contractility-deficient PGCs (cdPGCs). ns = non-significant (Mann-Whitney test).

(B) Rac1 FRET ratios for control PGCs or cdPGCs expressing either control or dominant-negative (DN) Rac1 protein. ****, P-value < 0.0001; ***, P-value = 0.0006 (Mann-Whitney test).

(C, D) Actin distribution in a control PGC (C) and a cdPGC (D), both expressing DN Rac1 (Lifeact, green, Video S7), with the respective graphs showing the average polarity value of actin over time (see STAR Methods). Shaded areas represent SEM.

(E, F) Actin distribution in a control PGC (E) or a cdPGC (F), both expressing constitutively active (CA) Rac1 (Lifeact, green, Video S8) with the graph showing the average polarity value of actin over time. Shaded areas represent SEM. Scale bars: 10 µm. The dashed lines represent the threshold set between apolar tumble phases (<0.1) and polar run phases (>0.1). n is the number of cells analyzed.

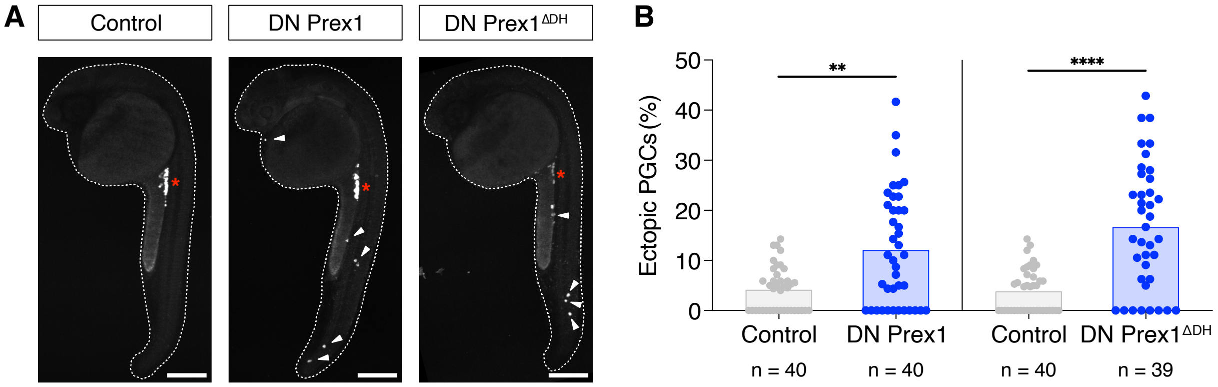


**Figure S3. Knockdown of Prex1 impairs PGC arrival at the gonad region, Related to Figure 3**

(A) 24 hpf embryos with GFP-labeled PGCs that express either control, DN Prex1 or DN Prex1$^{\Delta DH}$ protein. Red Asterisks point at the migration target region and white arrowheads at ectopic cells. Scale bar: 200 µm.

(B) The fraction of ectopic PGCs upon expression of either control, DN Prex1 and Prex1 ΔDH proteins. **, P-value = 0.0022; ****, P-value < 0.0001 (Mann-Whitney test). n is the number of cells analyzed.

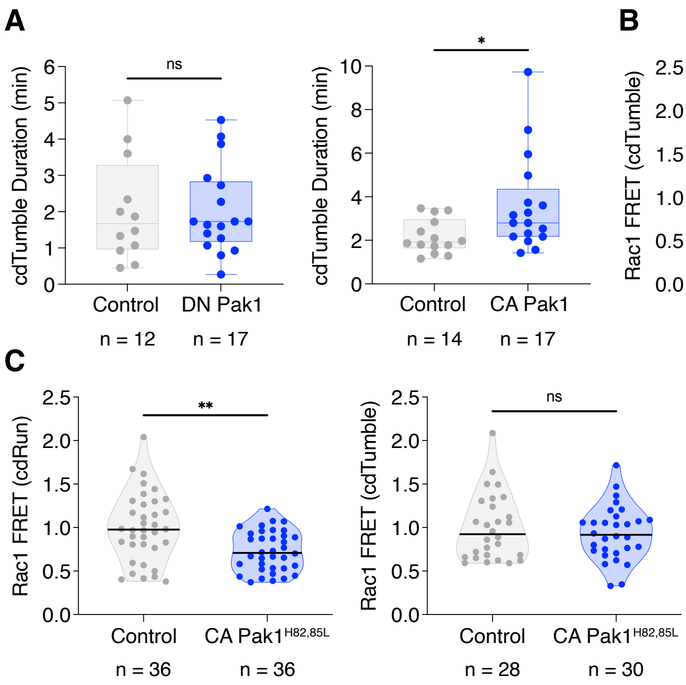


**Figure S4. Effect of Pak1 activity on the duration of tumble phases and on Rac1 activity, Related to Figure 4**

(A) Duration of contractility-deficient (cd)Tumble phases in control cdPGCs and cdPGCs expressing DN Pak1 (left graph) and in control cdPGCs and cdPGCs expressing CA Pak1 (right graph). ns = non-significant; *, P-value = 0.0227 (Mann-Whitney test).

(B) Rac1 FRET ratios during cdTumble phases in control cdPGCs and in cdPGCs expressing DN Pak1 (left graph) and in control cdPGCs and cdPGCs expressing CA Pak1 (right graph). ns = non-significant; *, P-value = 0.0423 (Mann-Whitney test).

(C) Rac1 FRET ratios measured in control cdPGCs and in cdPGCs expressing CA Pak1$^{H82,85L}$ during cdRun or cdTumble phases. **, P-value = 0.0018; ns = non-significant (Mann-Whitney test). n is the number of cells analyzed.

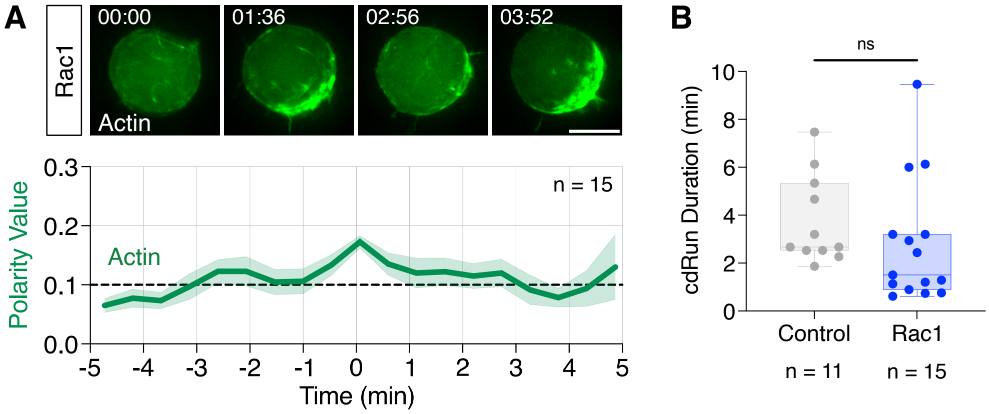


**Figure S5. Actin dynamics and run-phase duration in cdPGCs overexpressing Rac1, Related to Figure 4**

(A) Dynamics of actin distribution (Lifeact, green) in a contractility-deficient PGC (cdPGC) overexpressing Rac1 (Video S15). The graph presents the average polarity value of actin over time (see STAR Methods).

(B) Duration of cdRun phases of control cdPGCs and cdPGCs overexpressing Rac1. ns = non-significant (Mann-Whitney test). Shaded areas represent SEM. Scale bars: 10 µm. The dashed lines represent the threshold set between the apolar tumble phases (<0.1) and polar run phases (>0.1). n is the number of cells analyzed.

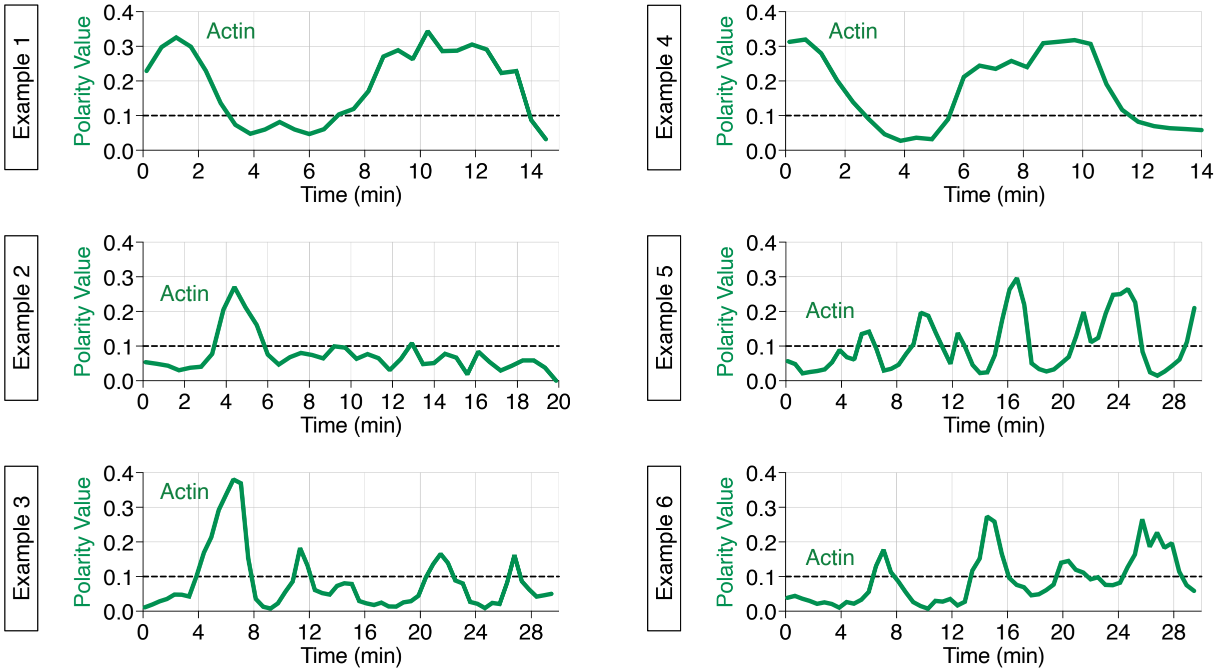


**Figure S6. Examples for actin dynamics in individual cdPGCs expressing CA Pak1, Related to Figures 4 and 6**

The graphs present the polarity value of actin over time of an individual cdPGC expressing CA Pak1. The dashed lines represent the threshold set between apolar tumble phases (<0.1) and polar run phases (>0.1).

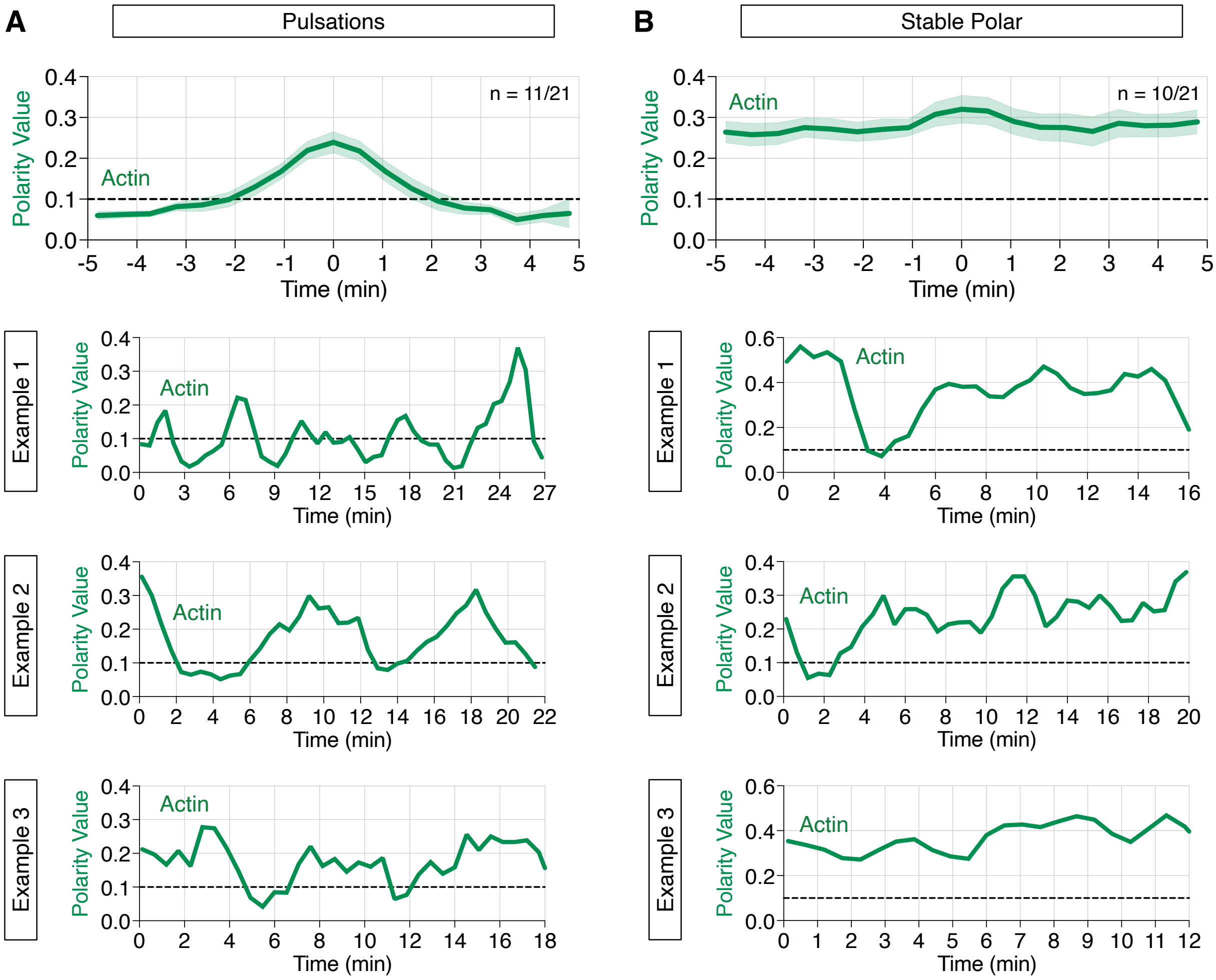


**Figure S7. Actin dynamics of pulsatory or stable polar cdPGCs expressing DN Pak1, Related to Figures 4 and 6**

(A) The upper graph presents the average polarity value of actin over time (see STAR Methods) in cdPGCs expressing DN Pak1, which show pulsatory behavior (run phases <10 min). The lower graphs show individual examples of the polarity value of actin over time in cdPGCs.

(B) The upper graph presents the average polarity value of actin over time in cdPGCs expressing DN Pak1, which show stable polar behavior (run phases >10 min). The lower graphs show examples of the polarity value of actin over time in individual cdPGCs. The dashed lines represent the threshold set between apolar tumble phases (<0.1) and polar run phases (>0.1). n is the number of cells analyzed.

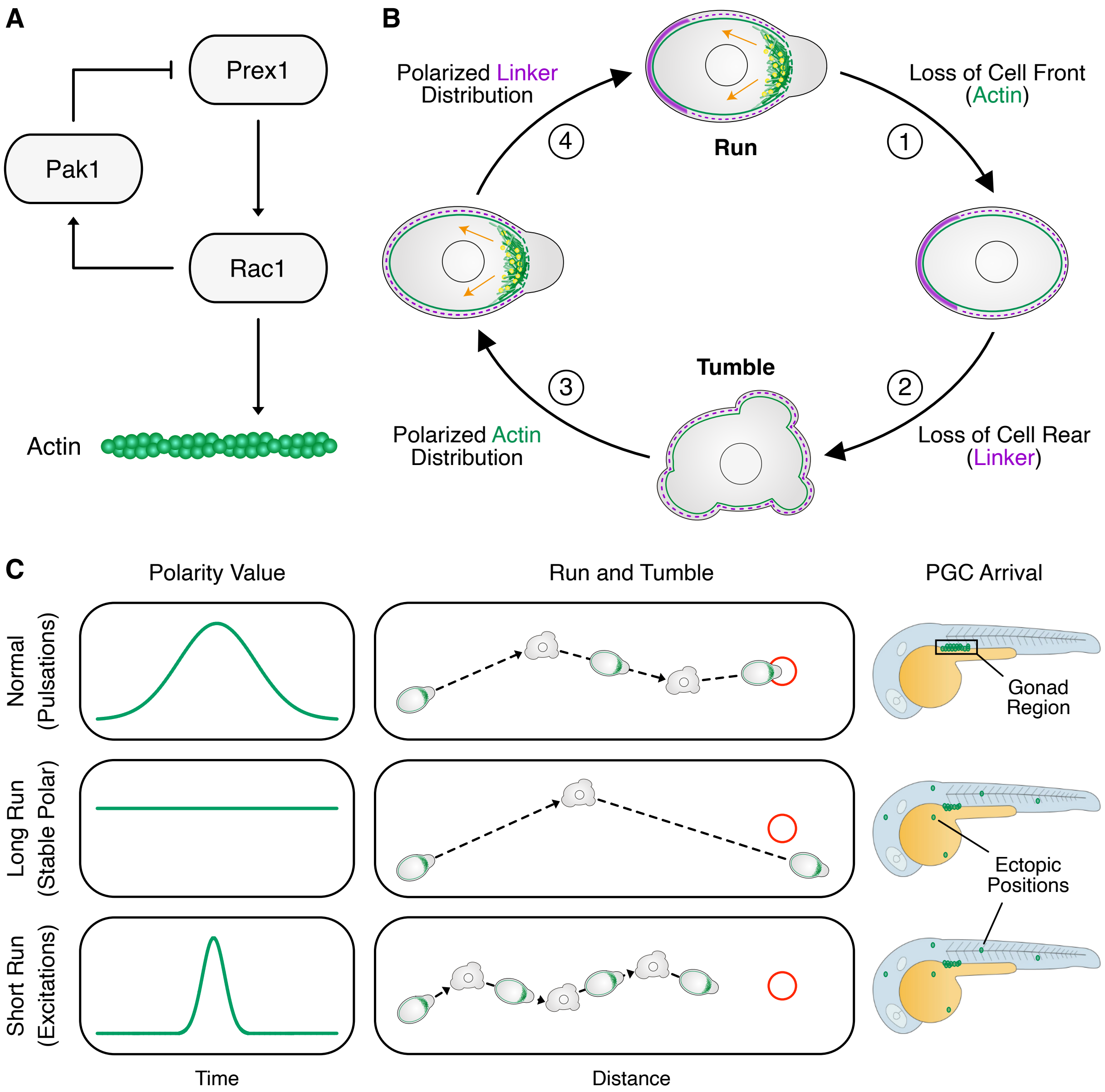


**Figure S8. A model for polar pulsations and their function in PGCs, Related to Discussion**

(A) A schematic representation of the molecular network that controls the polar pulsations in PGCs. Pointed arrows (→) indicate activation, and blunt arrows (–I) indicate inhibition. F-actin is presented as chains of green spheres.

(B) Illustration of the steps involved in the transition between run and tumble phases. (1) The initial step in the transition from run to tumble is the loss of the cell front, as can be evaluated by loss of polarized actin (green) distribution due to inhibition of Prex1 activity by Pak1. The depolymerization of actomyosin abrogates retrograde flow (orange arrows), which disrupts the shuttling of linker proteins such as Ezrin and Esyt2a to the cell rear (2). Under these conditions, both the front and the rear of the cells are not defined, and the cell enters the tumble phase. (3) The establishment of polarity during the tumble to run transition can be initiated by polarized activation of Rac1. The activation of this cascade can either be stochastic or biased by the activation of Cxcr4b by its ligand Cxcl12a. Actomyosin-based contractility then generates retrograde flow that (4) leads to the transport of linker proteins to the cell rear.

(C) Illustration showing the effect of different pulsation frequencies (left panels) on run and tumble behavior (middle panels) and PGC arrival at the migration target region (right panels). The dashed lines label the migration path of the cells. The red circle represents the migration target (i.e. the region where the gonad develops).

**Video S1. Dynamics of actin distribution during run and tumble phases, Related to Figure 1**
A representative time-lapse movie showing the dynamics of actin distribution (Lifeact, green) in a migrating PGC. The graph shows the polarity value of actin over time. Time in mm:ss format. Scale bar: 10 µm.

**Video S2. Dynamics of actin and Rac1 GTP distribution during run to tumble transition, Related to Figure 1**
A representative time-lapse movie showing the dynamics of actin (Lifeact, green) and GTP-bound Rac1 (Pak1 GBD, red) distribution in a migrating PGC during the run to tumble transition. The graph shows the polarity values of actin (green) and those of GTP-bound Rac1 (red) over time. Time in mm:ss format. Scale bar: 10 µm.

**Video S3. Dynamics of actin and Esyt2a distribution during run to tumble transition, Related to Figure 1**
A representative time-lapse movie showing the dynamics of actin (Lifeact, green) and that of extended synaptotagmin-like 2a (Esyt2a, magenta) distribution in a migrating PGC during the run to tumble transition. The graph shows the polarity values of actin (green) and Esyt2a (magenta) over time. Time in mm:ss format. Scale bar: 10 µm.

**Video S4. Polar pulsations of actin in a cdPGC, Related to Figure 2**
A representative live-cell time-lapse movie showing the dynamics of actin distribution (Lifeact, green) in a contractility-deficient PGC (cdPGC) during the transition from cdRun and cdTumble. The graph shows the polarity value of actin over time. Time in mm:ss format. Scale bar: 10 µm.

**Video S5. Dynamics of Ezrin distribution during polar pulsations of actin in a cdPGC, Related to Figure 2**
A representative time-lapse movie showing the dynamics of actin (Lifeact, green) and Ezrin (magenta) distribution in a contractility-deficient PGC (cdPGC) during transition between cdRun and cdTumble . The graph shows the polarity values of actin (green) and Ezrin (magenta) over time. Time in mm:ss format. Scale bar: 10 µm.

**Video S6. Polar pulsations of actin and Rac1 GTP in a cdPGC, related to Figure 2**
A representative time-lapse movie showing the dynamics of actin (Lifeact, green) and GTP-bound Rac1 (Pak1 GBD, red) distribution in a contractility-deficient PGC (cdPGC) during the transition between cdRun and cdTumble. The graph shows the polarity values of actin (green) and GTP-bound Rac1 (red) over time. Time in mm:ss format. Scale bar: 10 µm.

**Video S7. Dynamics of actin distribution in a PGC and a cdPGC expressing DN Rac1, Related to Figure S2**
A representative time-lapse movie showing the dynamics of actin (Lifeact, green) in a PGC expressing dominant-negative (DN) (1$^{st}$ segment) and in a contractility-deficient PGC (cdPGC) expressing DN Rac1 (2$^{nd}$ segment). The graphs show the polarity value of actin over time for both cases. Time in mm:ss format. The dashed lines represent the threshold set between apolar tumble phases (<0.1) and polar run phases (>0.1). Scale bar: 10 µm.

**Video S8. Dynamics of actin distribution in a PGC and a cdPGC expressing CA Rac1, Related to Figure S2**

A representative time-lapse movie showing the dynamics of actin distribution (Lifeact, green) in a PGC expressing constitutively active Rac1 version ( CA Rac1, 1st segment) and in a contractility-deficient PGC (cdPGC) expressing CA Rac1. The graphs show the polarity value of actin over time. Time in mm:ss format. The dashed lines represent the threshold set between apolar tumble phases (<0.1) and polar run phases (>0.1). Scale bar: 10 µm.

**Video S9. Dynamics of actin distribution in a cdPGC expressing DN Prex1, Related to Figure 3**

A representative time-lapse movie showing the dynamics of actin distribution (Lifeact, green) of a contractility-deficient PGC (cdPGC) expressing a dominant-negative form of the Prex1 protein (DN Prex1). The graph shows the polarity value of actin over time. Time in mm:ss format. The dashed line represents the threshold set between apolar tumble phases (<0.1) and polar run phases (>0.1). Scale bar: 10 µm.

**Video S10. Dynamics of actin distribution in a cdPGC overexpressing Prex1, Related to Figure 3**

A representative time-lapse movie showing the dynamics of actin distribution (Lifeact, green) in a contractility-deficient PGC (cdPGC) overexpressing Prex1. The graph shows the polarity value of actin over time. Time in mm:ss format. The dashed line represents the threshold set between apolar tumble phases (<0.1) and polar run phases (>0.1). Scale bar: 10 µm.

**Video S11. Dynamics of actin distribution in a cdPGC expressing CA Prex1, Related to Figure 3**

A representative time-lapse movie showing the dynamics of actin distribution (Lifeact, green) in a contractility-deficient PGC (cdPGC) expressing the constitutively active form of the Prex1 protein (CA) Prex1). The graph shows the polarity value of actin over time. Time in mm:ss format. The dashed line represents the threshold set between apolar tumble phases (<0.1) and polar run phases (>0.1). Scale bar: 10 µm.

**Video S12. Dynamics of actin distribution in a cdPGC expressing DN Pak1, Related to Figure 4**

A representative time-lapse movie showing the dynamics of actin distribution (Lifeact, green) in a contractility-deficient PGC (cdPGC) expressing dominant-negative form of the Pak1 protein (DN) Pak1. The graph shows the polarity value of actin over time. Top left corner shows the time in mm:ss format. The dashed line represents the threshold set between apolar tumble phases (<0.1) and polar run phases (>0.1). Scale bar: 10 µm.

**Video S13. Dynamics of actin distribution in a cdPGC expressing CA Pak1, Related to Figure 4**

A representative time-lapse movie showing the dynamics of actin distribution (Lifeact, green) in a contractility-deficient PGC (cdPGC) expressing constitutively active form of the Pak1 protein (CA Pak1). The graph shows the polarity value of actin over time. Time in mm:ss format. The dashed line represents the threshold set between apolar tumble phases (<0.1) and polar run phases (>0.1). Scale bar: 10 µm.

**Video S14. Dynamics of actin distribution in a cdPGC expressing Rac1C40, related to Figure 4**

A representative time-lapse movie showing the dynamics of actin distribution (Lifeact, green) in a contractility-deficient PGC (cdPGC) expressing Rac1$^{C40}$ protein. The graph shows the polarity value of actin over time. Time in mm:ss format. The dashed line represents the threshold set between apolar tumble phases (<0.1) and polar run phases (>0.1). Scale bar: 10 µm.

**Video S15. Dynamics of actin distribution in a cdPGC overexpressing Rac1, related to Figure S5**

A representative time-lapse movie showing the dynamics of actin distribution (Lifeact, green) in a contractility-deficient PGC (cdPGC) overexpressing Rac1. The graph shows the polarity value of actin over time. Time in mm:ss format. The dashed line represents the threshold set between apolar tumble phases (<0.1) and polar run phases (>0.1). Scale bar: 10 µm.
